# N-Myc downstream regulated gene 1b is a regulator of cell adhesion during early muscle development

**DOI:** 10.1101/2025.05.08.652902

**Authors:** P.K. Chowdhary, R.M. Brewster

## Abstract

The complexity of cellular functions necessitates intricate pathways that govern protein synthesis, transport, and degradation; yet the mechanisms governing the trafficking of key developmental proteins *in vivo* remain incompletely understood. N-Myc downstream-regulated genes (NDRGs) have recently emerged as stress-responsive proteins with roles in protein trafficking, but their functions during normal embryogenesis are largely unknown. Here, we identify N-Myc downstream-regulated gene 1b (Ndrg1b) as a novel regulator of N-cadherin (N-cad, *cdh2*) trafficking during zebrafish muscle development. We show that *ndrg1b* is broadly expressed during embryonic and larval stages and is required for proper skeletal muscle morphogenesis. Loss of Ndrg1b disrupts N-cad localization and impairs cell adhesion *in vitro*, highlighting a key role in maintaining tissue integrity. Mechanistically, Ndrg1b promotes the recycling of N-cad to the plasma membrane, identifying it as a novel component of the endocytic trafficking machinery *in vivo*. Given N-cad’s central role in development and disease, these findings provide new insight into how its localization is fine-tuned during morphogenesis. This work also expands the functional repertoire of NDRGs beyond stress response, suggesting that Ndrg1b – and potentially other NDRG family members – may broadly regulate transmembrane protein trafficking in a context-dependent manner. Furthermore, these findings may shed greater light on the etiology of Charcot-Marie-Tooth Disease type 4D, which has been linked to mutations in NDRG1.

**GRAPHICAL ABSTRACT:** (generated in Biorender)

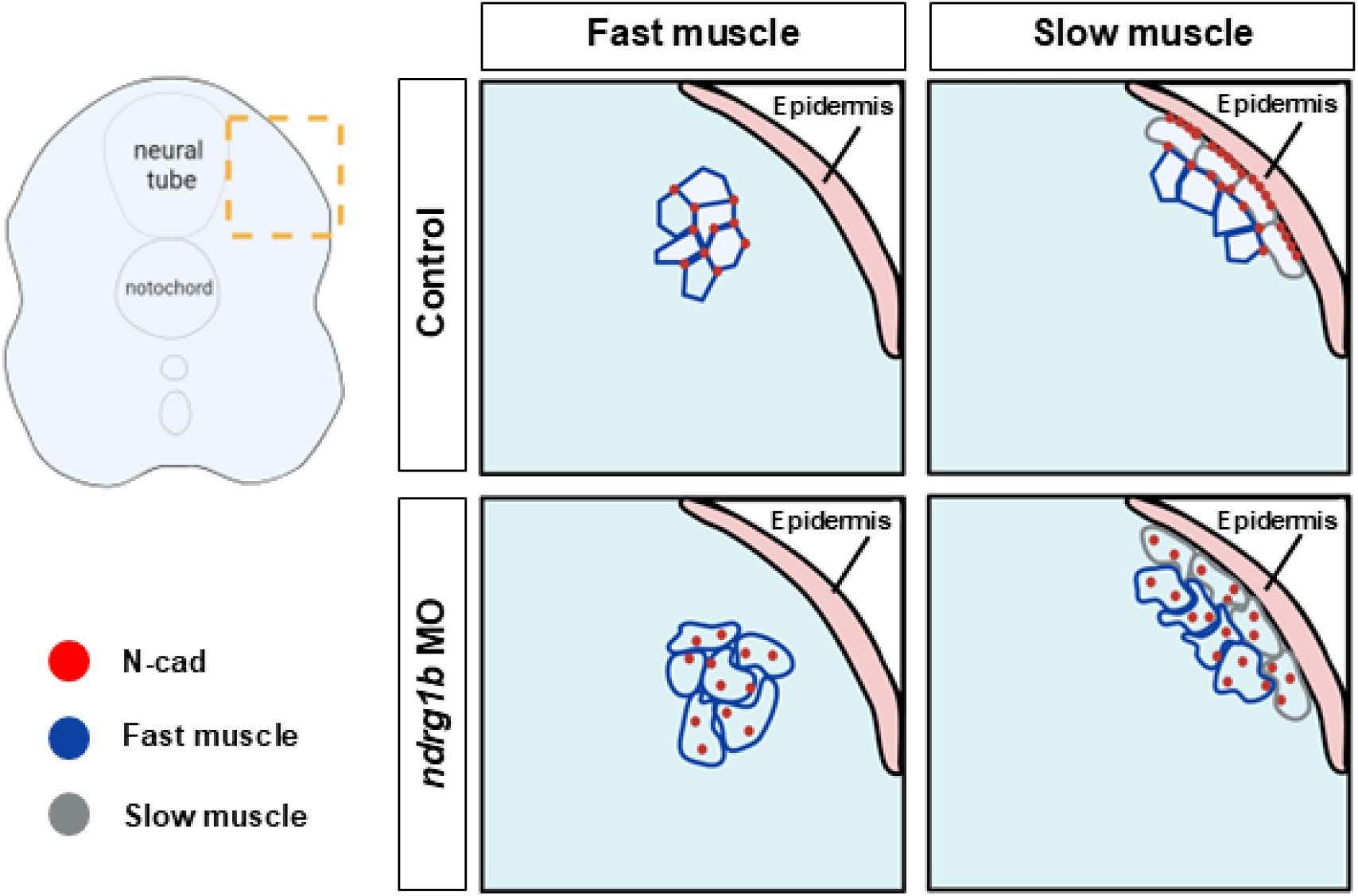

**Key takeaways:**

- Zebrafish *ndrg1b* is ubiquitously expressed in early development
- Loss of Ndrg1b is strikingly similar to the loss of N-cadherin
- Ndrg1b is required for the proper subcellular localization of N-cadherin in muscle cells and for the proper morphogenesis of this organ
- Loss of Ndrg1b reduces the ability of cells to aggregate *in vitro*

## Introduction

NDRGs (N-Myc Downstream Regulated Genes) are members of the α/β hydrolase family that are catalytically inactive yet still serve many broad functions in cell proliferation, signaling, and differentiation. Recent studies on such “pseudoenzymes” or “dead enzymes” – catalytically inactive enzyme homologs – reveal that such proteins can, in fact, fulfill a broad range of biochemical roles (Adrain, 2020). Owing to these capabilities, there is a growing need to identify and investigate such proteins as they may serve as versatile biochemical regulators in numerous cellular processes. The first family member, NDRG1, was originally identified as a target repressed by proto-oncogenes MYC and N-MYC (Shimono et al. 1999) and has since been studied under various aliases (e.g., RTP, Rit42, Cap43, PROXY1) in the contexts of cancer, hypoxia, and peripheral neuropathy (Kokame et al., 1996; Zhou et al., 1998; Piquemal et al.,1998;Echaniz-Laguna et al., 2007; Kachhap et al., 2007; Kitowska and Pawelczyk, 2010; Park et al., 2022).

As NDRGs encode enzymatically inactive hydrolases, their molecular functions remain poorly understood. Emerging evidence suggests that they may act as adapter proteins, serving as molecular scaffolds that facilitate interactions, complex formation, or signaling by binding with multiple partners. There is also growing evidence that adapter proteins regulate the trafficking of key proteins by recruiting cargo to vesicle coat proteins and by linking vesicles to motor proteins involved in intracellular transport (Park and Guo, 2014). Unlike most adapter proteins that act in specific trafficking steps, NDRGs have been linked to a broad range of trafficking processes – including endocytosis, exocytosis, and recycling – placing them at the intersection of multiple membrane transport pathways (Sugiki et al., 2004; Hunter et al., 2005; Tu et al., 2007; Pietiäinen et al., 2013; Mustonen et al., 2021). However, virtually all studies to date have examined these roles in cultured cells and/or under stress conditions, such as disease, hypoxia, or water immersion stress (Kachhap et al., 2007; Miyata et al., 2011; Park et al., 2022). Whether NDRGs regulate protein trafficking during normal development *in vivo* remains largely unexplored.

Our lab has previously demonstrated that zebrafish Ndrg1a plays a role in energy conservation under hypoxia by downregulating Na+/K+ ATPase in the kidney (Park et al., 2022). With zebrafish expressing two paralogs of NDRG1 – *ndrg1a* and *ndrg1b* – there is an urgent need to investigate *ndrg1b*, the functions of which remain largely unknown. While some studies have reported restricted expression of *ndrg1b* in the eye and epiphysis (Thisse et al., 2001; Tovin et al., 2012; Takita et al., 2016; Le et al., 2021), we find that its expression is much broader – consistent with Li et al. (2016) and more analogous to human NDRG1 and Xenopus *ndrg1*, rather than the localized expression of *ndrg1a* in the kidney and intestine (Thisse et al., 2001; Kyuno et al., 2003).

Here, we investigate Ndrg1b’s function during zebrafish embryonic development, focusing on its role in trafficking of the cell adhesion molecule N-cadherin (N-cad, *cdh2*). Given the stringently organized distribution of cadherins in the muscle, as well as the well-characterized role of N-cad in muscle migration and fusion, we focused on this tissue as a model for how Ndrg1b may regulate N-cad. The well-studied defects of N-cad in muscle migration and fusion allow for a strong readout of Ndrg1b’s effects in this tissue. We show that *ndrg1b*- and *N-cad*-depleted embryos present similar morphological defects and that their transcript distributions overlap both temporally and spatially. Loss of Ndrg1b reduces total N-cad levels and causes accumulation of intracellular N-cad puncta in muscle, consistent with impaired recycling to the plasma membrane. These findings are supported by *in vitro* evidence that loss of Ndrg1b disrupts cell adhesion.

This work is among the first to demonstrate a developmentally essential, stress-independent role for an NDRG family member in regulating protein trafficking *in vivo*. It identifies Ndrg1b as a novel adapter protein that fine-tunes N-cad localization during muscle development, thereby expanding the known repertoire of NDRG functions and revealing a new avenue for fine-tuning cell adhesion in the context of early embryonic development. The significance of this work is underscored by the well-documented role of N-cad in early development and cancer metastasis and may deeper inform our understanding of Charcot-Marie-Tooth disease type 4D, a neuropathy which has been linked to mutations in NDRG1.

## Results

### Relationship among NDRGs

Most organisms have four NDRGs, named NDRG[1–4]; however, owing to a genome duplication event in the teleost lineage ∼320 million years ago, many teleost genes have multiple paralogs of their mammalian counterparts (Fig. 1a). Zebrafish and other teleosts such as the killifish, channel catfish, and common carp have NDRG paralogs Ndrg1a and Ndrg1b compared to human NDRG1, as well as Ndrg3a and Ndrg3b compared to human NDRG3. The different NDRG paralogs share approximately 53-65% homology at the protein level, yet respective homologs exhibit higher identity between different organisms. All zebrafish paralogs of Ndrg1 diverge most from the NDRG1 protein sequence in the N-terminus; however, Ndrg1b contains two deletions in the C-terminus and is lacking three repeats of a 10-a.a. sequence that is conserved among the others. In other teleosts with paralogs of NDRG1, the Ndrg1b ortholog also lacks this C-terminus three-peat (Fig. 1b,c). Importantly, phylogenetic analysis among these NDRG members does not reveal a clear evolutionary relationship. This ambiguity is further supported by gene synteny analysis of the genetic neighbors of the zebrafish Ndrgs – *ndrg1a* shares 8 neighbors within the surrounding 50 genes with human *ndrg1* whereas *ndrg1b* shares 9 (Fig. 1d).

**Figure 1.**
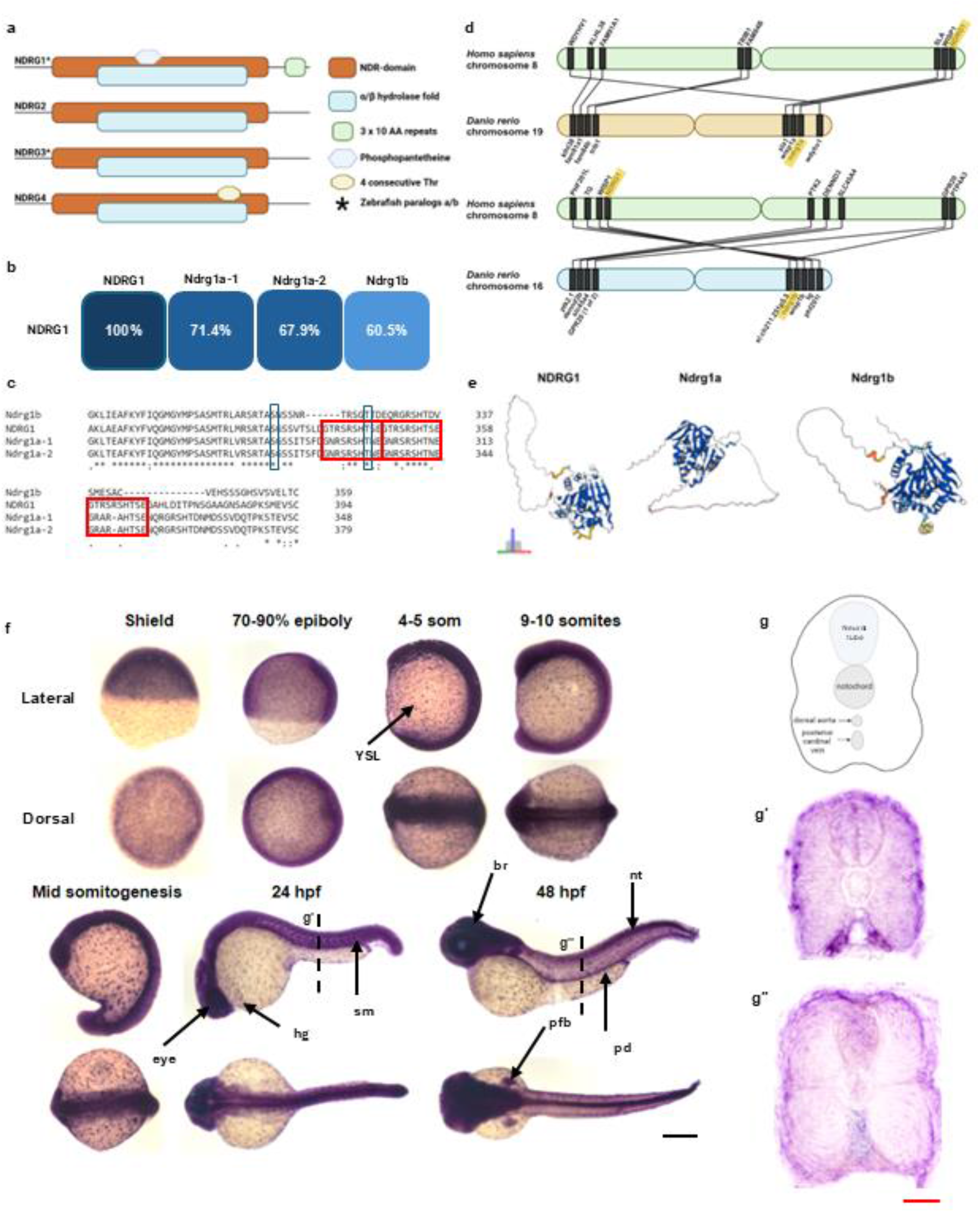
NDRG family of proteins and zebrafish *ndrg1b* expression. (a). NDRG members consist of NDRG 1–4, sharing 57-68% amino acid identity. Zebrafish Ndrgs consist of further paralogs Ndrg1a/1b and Ndrg3a/3b. All NDRG members contain the NDR domain and the α/β hydrolase fold. Adapted from Melotte et al., 2010 (permission obtained). (b) ClustalW alignment of zebrafish Ndrg1a-1, Ndrg1a-2, Ndrg1b, and human NDRG1 and their overall sequence similarity summarized. (c) C-terminal tandem repeats present in NDRG1, NDRG1a-1, and NDRG1a-2. (d) Gene synteny analysis of nearest 50 neighbors comparing *ndrg1a* to *ndrg1* (top) and *ndrg1b* to *ndrg1* (bottom). (e) AlphaFold predicted structures for NDRG1, Ndrg1a, and Ndrg1b. The α/β hydrolase fold is conserved amongst all three, and the greatest differences in structure arise from the intrinsically disordered C-terminus. (f) Use of mRNA *in situ* hybridization to investigate the levels and localization of *ndrg1b* throughout early development. In early development, *ndrg1b* mRNA is present ubiquitously in the embryo as well as in a punctate-like manner on the yolk syncytial layer (YSL). At 24 hpf, *ndrg1b* expression is strongest in the eye, but remains throughout the YSL, somites (sm), and hatching gland (hg). At 48 hpf, expression is strongest in the eye, brain (br), pectoral fin bud (pfb), pronephric duct (pd), and neural tube (nt). (g) Schematic of transverse section from somites. (g’,g’’) *ndrg1b* in the somites is localized superficially, just beneath the epidermis, and most lateral relative to the notochord and spinal cord. Schematics in a, d, and g generated in Biorender. Scale: black = 250 µm; red = 20 µm.

Although the crystal structures for zebrafish Ndrg1a and Ndrg1b have not yet been resolved, it is possible to predict 3D protein architecture through AI algorithms such as AlphaFold. Using these models can help guide predictions about how differences in structure may drive differences in function. The comparison among structures for NDRG1, Ndrg1a, and Ndrg1b is shown in Figure 1e (Jumper et al., 2021; Varadi et al., 2021). Notably, despite the greater difference in sequence homology between NDRG1 and Ndrg1b as well as the lack of the C-terminal three-peat in Ndrg1b, the C-terminal folding between NDRG1 and Ndrg1b is more similar in appearance than NDRG1 compared to Ndrg1a. The C-terminus of NDRG1 is more rod-shaped, which is mimicked in Ndrg1b, whereas the C-terminus of Ndrg1a is more hanger-shaped. Of note, however, the N- and C-termini of NDRG1 are intrinsically disordered regions lacking a stable conformation. While it is estimated that AlphaFold can identify conditionally folding intrinsically disordered regions (IDRs) at a precision rate as high as 88% with a 10% false positive rate, it is not possible to capture the transient nature of these structures and how they may change under varying conditions (Alderson et al., 2023; Wilson et al., 2022). It is additionally important to reiterate that the 3D structures of Ndrg1a and Ndrg1b have not been validated experimentally, and these models are theoretical. While sequence and structural similarities provide valuable information, greater insights into the relationship among NDRG orthologs may be gleaned by studying their expression and functions in different tissues.

### *ndrg1b* is broadly expressed during early development

The previous and limited studies on *ndrg1b* do not report on the dynamics of its expression throughout early embryonic development in zebrafish. We therefore first documented the expression of *ndrg1b* at earlier stages of development and found that *ndrg1b* is ubiquitously expressed throughout the embryo. Throughout mid-somitogenesis, *ndrg1b* is broadly expressed in the embryo and in a punctate-like manner in the yolk syncytial layer (YSL). At 24 hours post fertilization (hpf), expression is most prominent in the eye, central nervous system, and somites, whereas at 48 hpf expression is still broad but strongest throughout the brain and spinal cord (Fig. 1f). Cross sections through the trunk reveal that *ndrg1b* becomes more localized superficially, just deep to the epidermis, and laterally from the spinal cord and notochord (Fig. 1g). This pattern is consistent with the spatial organization of slow muscle fibers and the zone of N-cad expression in the myotome, which is further explored below. The ubiquitous expression of *ndrg1b* is more closely related to that of NDRG1, which is also reportedly expressed in many tissues (obtained from post-mortem adults). NDRG1 is also expressed in the human placenta, which is analogous to the zebrafish yolk where *ndrg1b* is expressed (Choi et al., 2007). In contrast, *ndrg1a* expression is more localized to the zebrafish kidney and ionocytes at these stages (Carithers et al., 2015; GTEx; Thisse et al., 2001).

### Loss of *ndrg1b* function results in severe morphological defects that resemble those of N-cad depleted embryos

To better understand the function of Ndrg1b, we performed several loss-of-function experiments, which revealed key morphological defects upon depletion of *ndrg1b*. Typically, 24 hpf embryos display a straight, elongated trail, a clear midbrain-hindbrain boundary, and shrunken yolks as the developing embryo uptakes its nutrients (Fig. 2a, left).The knockdown of *ndrg1b* in splice-and translation-blocked morphants, as well as F0 crispants, resulted in severe developmental defects in normoxic embryos. Embryos lacking *ndrg1b* display abnormal/necrotic brain anatomy and curved/kinked tails with posterior bulges – perhaps indicative of cell adhesion defects resulting in failed neural tube fusion and disruptions to convergent extension. Embryos also display enlarged yolks suggesting improper nutrient uptake, which is not the focus of this work but is consistent with the expression of *ndrg1b* in the yolk (Fig. 2a, right). These defects are characterized as mild, moderate, or severe and are quantified in Fig. 2b. The developmental defects observed with the loss of Ndrg1b correlates with the ubiquitous expression of the gene reported above. Interestingly, the developmental anomalies observed in *ndrg1b* morphants are strikingly similar to the phenotypes reported in *N-cad* mutants (Malicki et al., 2003; Lele et al., 2002; Hong & Brewster, 2006) and are recapitulated in this work in a direct comparison using N-cad morpholinos as previously described in Lele et al., 2002. Similarly, N-cad mutants and morphants display curved tails with posterior bulges, enlargement of the hindbrain ventricle, and disorganized brain anatomy (Fig. 2c). The *ndrg1b* splice-blocking morpholino was validated and is used in all subsequent experiments (SI1), and embryos with mild phenotypes were used in all experiments described further.

**Figure 2.**
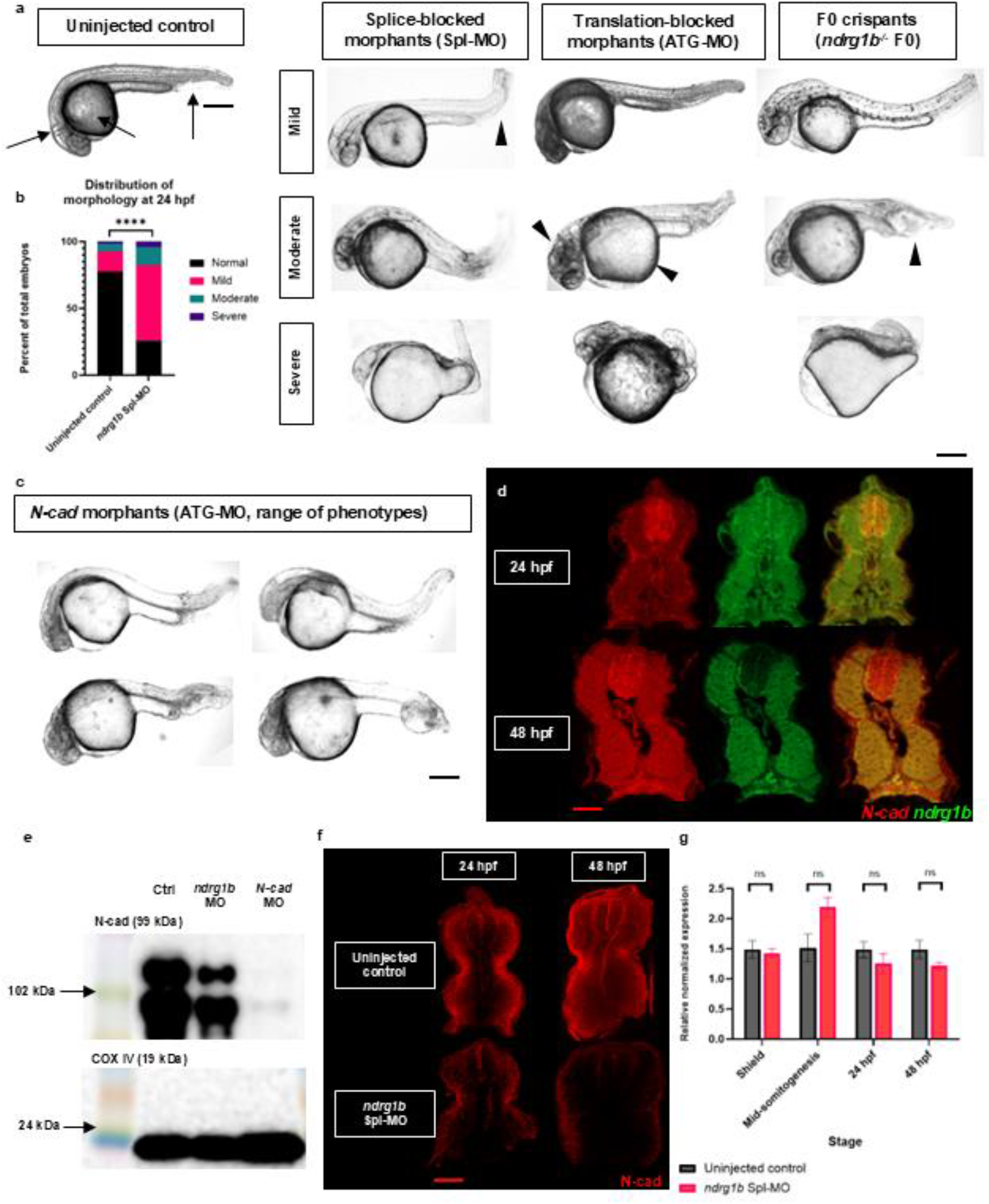
*ndrg1b* loss-of-function phenotype, mRNA overlap with *N-cad*, N-cad levels in *ndrg1b* morphants. (a) Multiple ndrg1b loss of function tools consistently result in disruptions to embryonic development. At 24 hpf, wild-type embryos (uninjected controls) have a straight body axis, 90° head-to-tail angle, distinct neural folds, and partially-depleted yolk sacs (black arrows). Splice-blocked morphants, translation-blocked morphants, and F0 crispants display severe abnormalities including enlarged yolks, abnormal neural folds, and curved/kinked tails with posterior bulges (arrowheads). (b) Quantification of morphology types observed in *ndrg1b* morphants (153 uninjected controls, 202 morphants analyzed). Fisher’s exact test performed for statistical analysis. (c) The range of phenotypes observed in *ndrg1b* knockdown embryos is consistent with those observed in *N-cad* morphants. (d) HCR-FISH reveals localization of *ndrg1b* and *N-cad* transcript in somites. Transverse sections reveal overlap in slow muscle cells and neural tube at 24 hpf, and throughout the muscle at 48 hpf. (e) *N-cad* transcript levels are not significantly different in *ndrg1b* morphants. Expression levels in qPCR were normalized to *ef1a*. Reactions were run in technical triplicate with 3 biological replicates of 3 pooled embryos (n=9). Bars represent standard error of the mean (SEM). Unpaired, two-tailed t-test used in statistical analysis. All fold changes were derived using the formula, 2^−(ΔΔ CT)^. (e) Western blot analysis showing partial loss of N-cad in *ndrg1b* morphants and validating complete loss of N-cad in *N-cad* morphants measured from total protein extracts in 24 hpf embryos. (f) Transverse sections of somites show (qualitatively) reduced levels and abnormal distribution of N-cad in *ndrg1b* morphants. Splice-blocking morpholino validation and lateral and dorsal views of wholemount HCR-FISH embryos throughout multiple stages available in Supplemental Information. Scale: black = 250 µm; red = 20 µm.

### *N-cad* and *ndrg1b* transcript are spatially overlapped in early development and in the somites

The striking similarity in the *ndrg1b* and *N-cad* loss-of-function phenotypes led us to postulate that perhaps Ndrg1b may regulate N-cad in some way. However, the spatial overlap of these genes has not been previously studied, which is crucial to their ability to interact. Although the effort to determine protein-protein colocalization or interactions are currently hindered by the lack of functional antibodies targeting zebrafish *ndrg1b*, the colocalization of *ndrg1b* and *N-cad* mRNA was tested using hybridization chain reaction (HCR). Through HCR we show that there is strong overlap of the expression domain of these two genes in the somites. Transverse sections of 24 hpf embryos show *N-cad* transcript enriched in slow muscle cells, though at 48 hpf this expression seems broad throughout the muscle cells. *ndrg1b* is broadly expressed throughout the muscle at 24 and 48 hpf. There is a strong overlap of transcript throughout the neural tube at 24 hpf, though *ndrg1b* appears basally enriched at 48 hpf (Fig. 2d; full views in SI2).

### Ndrg1b regulates the levels of N-cad protein, but not transcript

The similar loss-of-function phenotypes observed in both *ndrg1b* (Spl-MO) and *N-cad* (ATG-MO) morphants suggest that Ndrg1b may regulate N-cad. This hypothesis is further supported by previous evidence indicating that NDRGs function as regulators of protein trafficking and that NDRG1 specifically modulates E-cadherin recycling in prostate cancer cells (Kacchap et al., 2007). Previous work performed by Miyata et al. in 2011 also identifies a positive correlation between phosphorylated NDRG1 and adhesion molecules (β-catenin, α-catenin, and N-cadherin) in murine oligodendrocytes, though this relationship was only examined following water immersion restraint stress and did not study the role of NDRG1 in native conditions or through direct knockout. To test whether Ndrg1b controls N-cad levels, we examined N-cad levels in total protein extract from 24 hpf *ndrg1b* morphants using western blot analysis. The larger of the two bands detected by western blot likely corresponds to a post-translational modification, which, according to the size of the band shift, may be phosphorylation, palmitoylation, and/or glycosylation. Although these are not the focus of my work, it is interesting to note that both versions of N-cad are reduced in *ndrg1b* morphants relative to the control (Fig. 2e). As expected, N-cad protein levels in *N-cad* morphants are almost entirely diminished.

As protein extracts are obtained from whole embryos, it is uncertain whether these overall changes in N-cad expression are an accurate indication of the trend that follows in the somites specifically. While not quantitative, N-cad immunolabeled sections of 24 hpf and 48 hpf embryos do qualitatively suggest an overall decrease of N-cad protein in *ndrg1b* morphant somites (Fig. 2f). Zebrafish, like mammals, have two major muscle cell types – fast and slow twitch muscle fibers – and possess the experimental advantage of spatial segregation in the myotome as opposed to intermingled in mammals, thus making individual cell types easier to identify and study. Deep fast muscle cells predominantly express M-cadherin, while superficial slow muscle cells predominantly express N-cadherin. The homophilic attraction between like molecules of M-or N-cad, as well as cell-autonomous protrusive behaviors, drives the separation of slow muscle fibers laterally away from the neural tube and notochord, while fast muscle fibers remain deep and constitute the bulk of muscle mass (Granato, 2003; Cortés et al., 2003; Ono et al., 2015). Consistent with this, we observe that in wildtype embryos, N-cad is localized more superficially within the myotome (lateral relative to the notochord) and displays a gradient of expression where it is strongest in superficial slow muscle cells and weakest in deep muscle cells (Fig. 2f). *N-cad* mRNA levels were not significantly different between controls and *ndrg1b* morphants, suggesting that Ndrg1b regulates N-cad protein rather than mRNA levels, possibly by inhibiting N-cad degradation or by promoting its exocytosis and/or recycling (Fig. 2g). These possibilities are further explored in the discussion below.

### Ndrg1b controls N-cad subcellular distribution

Given the data above suggesting that Ndrg1b may regulate N-cad at the protein level, along with the breadth of previous research linking NDRGs to protein trafficking and the expression of adhesion molecules, we tested whether Ndrg1b regulates the subcellular localization of N-cad, perhaps resulting in reduced levels at the cell surface and thus causing the cell adhesion defects observed above. To visualize protein distribution in individual cells, we utilized membrane-targeted Enhanced Green Fluorescent Protein (mEGFP), which was injected as plasmid cDNA into the yolks of one-cell stage embryos and mosaically inherited throughout the embryo.

In both 24 and 48 hpf embryos – after slow muscle migration has taken place and the dynamics of N-cad turnover should be relatively unchanged – we observe that *ndrg1b* is required for the proper subcellular localization of N-cad. We observe that in fast muscle cells of control embryos, N-cad is strongly localized at the junctions that join cells in a parallel, tessellating array; in slow muscle cells, N-cad appears to be expressed throughout the lateral surface of the cell membrane (perhaps as an attachment to the overlying epidermis) and enriched at points of contact with other cells at the medial surface (Fig. 3).

**Figure 3.**
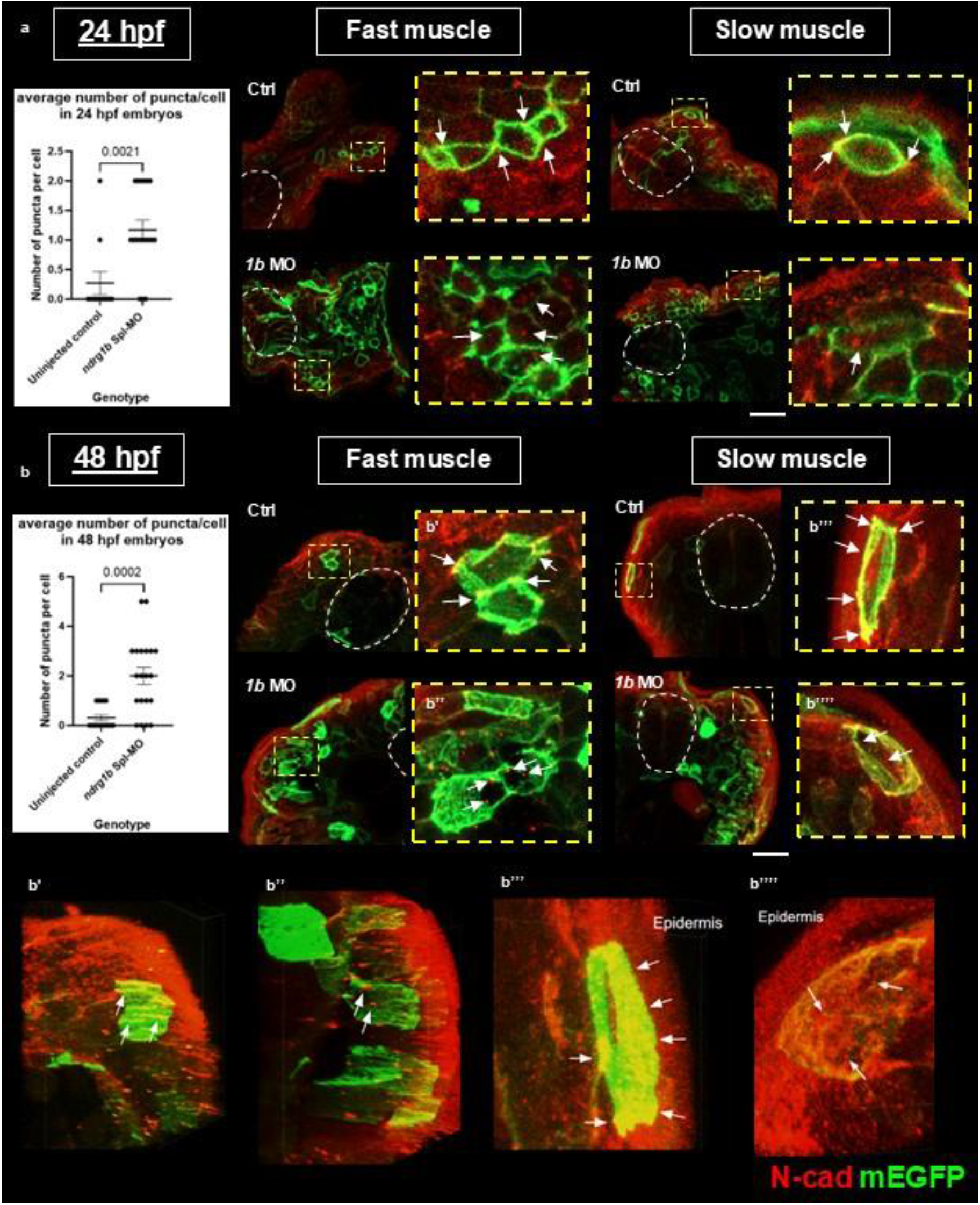
Loss of *ndrg1b* results in abnormal subcellular localization of N-cad in fast and slow muscle cells. Loss of *ndrg1b* causes altered subcellular localization of N-cad in fast and slow muscle cells in (a) 24 hpf and (b) 48 hpf embryos. In fast muscle cells of mEGFP-only control embryos, N-cad is localized to adherens junctions between cells, whereas in slow muscle cells N-cad is distributed across the membrane on the lateral surface and at attachment points on the medial surface. In mEGFP-*ndrg1b* morphants, this localization is disrupted and there is a significant increase in intracellular N-cad puncta. 24 hpf: average of 0.27 puncta/cell in wildtype, 1.16 puncta/cell in morphants. 48 hpf: average 0.31 puncta/cell in wildtype, 2 puncta/cell morphants. Unpaired, two-tailed t-tests used in statistical analyses. Bars represent standard error of the mean (SEM). Neural tube outlined in dotted white lines. (b’-b’’’’) Representative stills from 3D reconstructions (Supplemental Information) showing mislocalization of N-cad in fast and slow muscle cells of *ndrg1b* morphants. Individual channels for a and b are shown in Supplemental Information (SI3,4). Scale: white = 20 µm.

In *ndrg1b* morphants, both cell types show decreased levels of N-cad at the cell membrane and/or junctions and instead show an increase in the number of intracellular puncta (Fig. 3). In addition, *ndrg1b* morphants display aberrant muscle fiber organization as this tight array of pentagonal fibers is clearly disrupted in *ndrg1b* morphants and the plasma membrane of these cells appears more ruffled. Notably, a 3D reconstruction of fast and slow muscle cells in mEGFP-only controls (48 hpf) show the tight alignment of cells held together by cables of N-cad running along the length of the fibers at their junctions (Supplemental movies 1, 3; Fig 3b’, b’’’). Of note, the cables of N-cad that we observe in fast muscle fibers have not been previously reported to our knowledge. In mEGFP-*ndrg1b* morphants, this reconstruction not only highlights the increased levels of intracellular N-cad but also displays the very loose and disorganized arrangement of cells without proper junctions between muscle fibers (Supplemental movies 2, 4; Fig 3 b’’, b’’’’). Taken together, these results suggest a disruption to the trafficking of N-cad in *ndrg1b* morphants, which is explored further below.

### Ndrg1b mediates N-cad intracellular trafficking

Noting our observation that *ndrg1b* morphants display decreased levels of membrane N-cad and increased levels of intracellular N-cad, we postulated that the loss of Ndrg1b may result in a trafficking defect, consistent with those reported in the literature (Sugiki et al., 2004; Hunter et al., 2005; Tu et al., 2007; Kachhap et al., 2007; Mi et al., 2017; Mustonen et al., 2021). We therefore followed up on this finding by co-immunolabeling for N-cad with a marker for early endosomes (EEA1; early endosomal antigen one) and for recycling endosomes (Rab11) within the protein trafficking pathway. We find that N-cad appears to colocalize more strongly in EEA1-positive cells within the muscle domain in 24 and 48 hpf morphant embryos, suggesting its enrichment in early endosomes and improper intracellular trafficking (Fig. 4a,c; ROI for quantification in white box; example of EEA1-endosome noted with green arrow, N-cad-positive EEA1-endosome noted with yellow arrow). More specifically, we observe that in morphant embryos, N-cad is enriched in recycling endosomes (Fig 4b,c) but not lysosomes (LAMP1, SI7). These findings strongly support our working model that Ndrg1b regulates the intracellular trafficking of N-cad by promoting the endocytic recycling of N-cad to the cell surface.

**Figure 4.**
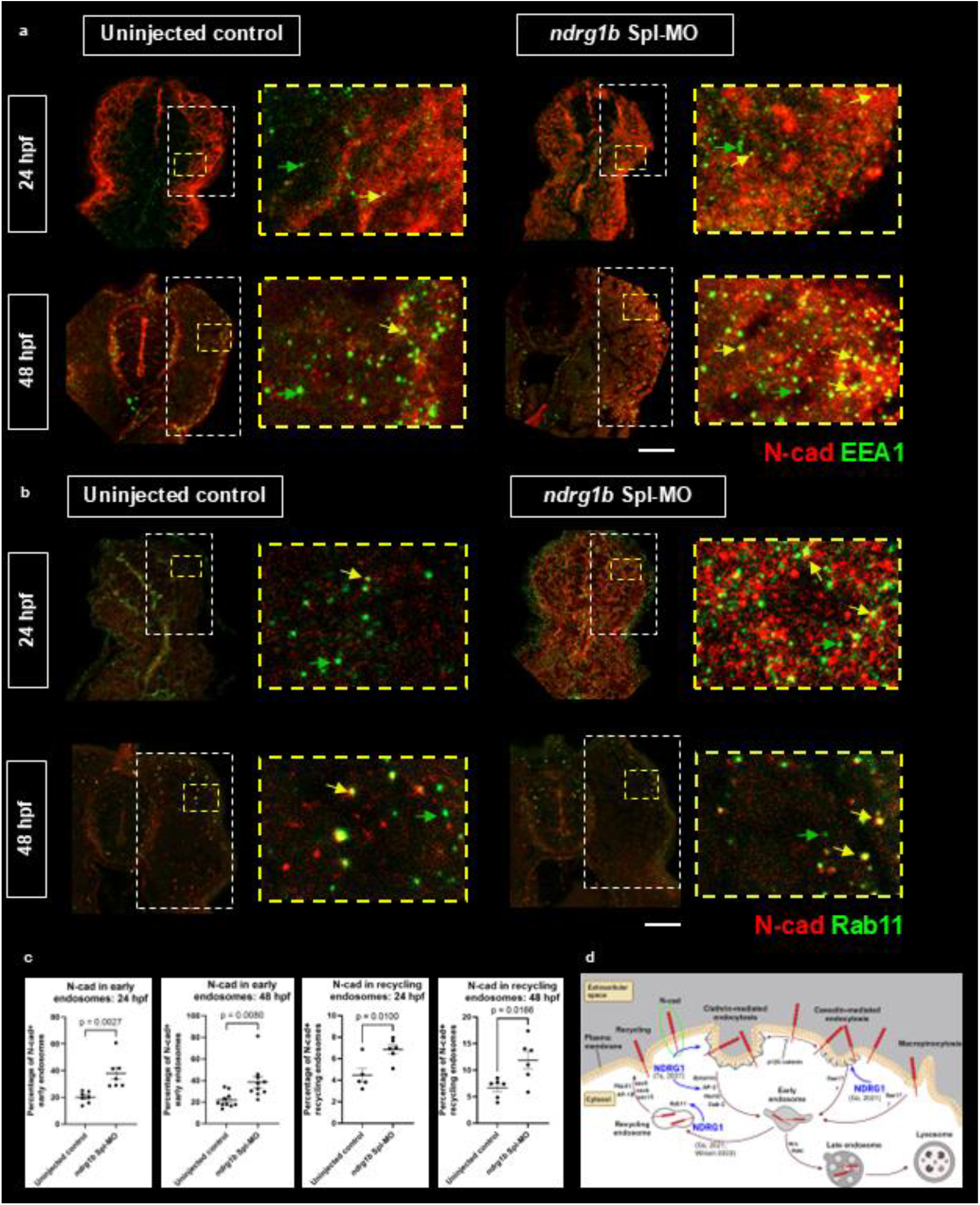
Enrichment of N-cad puncta in early and recycling endosomes. 24 and 48 hpf embryos co-labeled with N-cadherin (red) and either the early endosomal antigen 1 (a, EEA1, green), a marker for early endosomes, or Rab11 (b, green) for recycling endosomes. *ndrg1b* morphants show stronger colocalization of these markers, suggesting intracellular localization of N-cad and a hindrance in its trafficking pathway (green arrows showing labeled endosomes, yellow arrows showing N-cad+ endosomes). (c) Percentage of N-cad positive EEA1/Rab11 endosomes. 24 hpf EEA1: 20.11% N-cad+ controls, 37.70% N-cad+ morphants. 490 puncta quantified from 7 transverse sections across 3 control embryos; 533 puncta quantified from 7 transverse sections across 3 morphants. 48 hpf EEA1: 21.78% N-cad+ controls, 38.71% N-cad+ morphants. 1843 puncta quantified from 10 transverse sections across 5 control embryos; 1352 puncta quantified from 10 transverse sections across 5 morphants. 24 hpf Rab11: 4.45% N-cad+ controls, 6.86% N-cad+ morphants. 771 puncta quantified from 5 sections across 3 control embryos; 1570 puncta quantified from 6 sections across 3 morphants. 48 hpf Rab11: 6.66% N-cad+ controls, 11.87% N-cad+ morphants. 601 puncta quantified from 6 sections across 3 embryos; 679 puncta quantified from 6 sections across 3 morphants. Unpaired, two-tailed t-tests used for statistical analyses. Bars represent standard error of the mean (SEM). Individual channels for a and b, as well as data for LAMP1 labeled lysosomes, shown in Supplemental Information (SI5-7). (d) Trafficking pathway of N-cad and known binding partners of NDRG1. N-cad can be endocytosed through clathrin-mediated or caveolin-mediated endocytosis or through macropinocytosis before being sent to the lysosome for degradation or returned to the cell surface via recycling. Adapter molecules involved in these specific pathways are listed, NDRG1 binding partners are identified with blue arrows. Adapted from Kowalcyzk & Nanes, 2012 (permission obtained) and generated in Biorender. Scale: white = 20 µm.

The enrichment of N-cad in intracellular compartments of *ndrg1b* morphants may be contradictory with our earlier findings suggesting that N-cad levels are overall reduced with the loss of Ndrg1b. This may be an experimental artefact resulting from the inability to properly extract proteins that are sequestered within intracellular compartments, but it is possible that Ndrg1b may play a role in regulating the stability of N-cad as has been demonstrated for NDRG1 and the regulation of c-Myc (Wang et al., 2017; Deng & Richardson, 2023) or that improperly trafficked N-cad is subsequently marked for degradation. It is also possible that total N-cad levels are decreased, but specific tissues have different magnitudes of changes in N-cad levels. Future work will seek to address this through more stringent methods of protein extraction and by directly measuring the turnover of N-cad *in vivo*.

While NDRG1 has previously been linked to N-cad (Miyata et al., 2011), the work presented here provides the first mechanistic insight into how these two proteins may be linked. Although the trafficking pathway of N-cad has been well-studied, adapter proteins likely play a role in the fine-tuning of N-cad function and dynamic demands for cell adhesion. The trafficking pathway of N-cad is illustrated in Figure 4d, along with the identification of molecules within the pathway to which NDRG1 has been shown to bind directly. To date, there is no direct binding evidence between NDRG1 and N-cad, though it is likely that the protein trafficking abilities of NDRGs are attributed to its scaffolding function linking trafficking effectors and cargo.

### Ndrg1b regulates slow muscle migration

Many studies have established that the correct levels, distribution, and subcellular localization of N-cad are necessary for the proper development and organization of many different organs, including muscle tissue. The work described above has strongly established Ndrg1b as a novel regulator of N-cad transport in muscle tissue, and, as such, suggests that the loss of this protein will disrupt muscle development in an N-cad-dependent manner. The well-understood role of N-cad in muscle fiber migration and fusion allows for this tissue to be easily studied in the context of disrupted protein transport.

Currently, much of our understanding of NDRGs’ functions as regulators of protein trafficking comes from studies in cell culture, disease contexts, and/or cell stress. Our work addresses this gap by investigating the function of Ndrg1b in a developing organism and thereby increasing understanding of its relevance to cellular and physiological processes. One such process in which N-cad functions is in muscle development, where the differential expression of N-cad in muscle cells is critical for muscle migration and organization, and, at a cellular level, cell fate specification, migration, and fusion (Hollnagel et al., 2002; Demonbreun et al., 2015). In zebrafish, the spatial segregation of different muscle fibers is in part driven and maintained by the relative levels of M-and N-cad in specific cell types. During this cell migration, the sorting of M-and N-cad-expressing cells involves controlling the intracellular transport of these adhesion molecules as the cells slide past one another (Cortés et al., 2003; Granato et al., 2003). Although less well understood and not the focus of this paper, the turnover of cadherins in mature muscle fibers is also thought to play a role in signal transduction, maintaining muscle integrity, and adaptation to stress.

Given that N-cad is not properly localized and regulated in *ndrg1b* morphants, we speculated that developmental processes and functions that are dependent on the tight regulation of cell adhesion will be disrupted in *ndrg1b* morphants. We postulated that improper trafficking of N-cad in *ndrg1b* morphants results in a loss of cell adhesion that translates into defects in muscle development. In zebrafish, where fast and slow muscle fibers are spatially segregated, the differential expression and the precise balance of cell adhesion molecules is required to ensure the correct organization of this tissue resulting in superficial slow muscle fibers and deep fast muscle fibers. Slow muscle migration begins around 17 hpf in embryos and is complete by 24 hpf (Keenan et al., 2019).

The disruption of N-cad trafficking in *ndrg1b* morphants suggests that these embryos would display defects in cell adhesion and in processes that are dependent on N-cad function, such as slow muscle migration (Granato, 2003; Cortés et al., 2003). To test for defects in muscle migration, we immunolabeled embryos with the slow muscle marker F59 (recognizing myosin heavy chain 1) and imaged transverse sections to track the migration of individual fibers in embryos ranging from 17–48 hpf. In uninjected controls, slow muscle cells successfully migrate outward towards the periphery of the myotome (Fig. 5a, top; Fig. 5b). However, with the loss of *ndrg1b*, muscle fibers are severely disorganized as they do not completely migrate through the fast muscle domain (Fig. 5a, middle; Fig. 5b). Importantly, this defect persists, and is therefore not the result of a delay, shown prominently in 30 hpf and 48 hpf embryos that still display slow muscle fibers that have not fully migrated. Of note, the morphants selected for this analysis were those with mild anatomical defects, and as such the anomalies observed are not an indirect consequence of some larger disruption in tissue morphogenesis and organization. These defects observed in *ndrg1b* morphants are consistent with those observed in *N-cad* morphants (Fig. 5a, bottom; Fig. 5b), suggesting that *ndrg1b* regulates slow muscle migration in an N-cad-dependent manner. The defects in *N-cad* morphants appear more severe, which is unsurprising as losing the adhesion molecule itself would be more impactful than losing its regulator.

**Figure 5.**
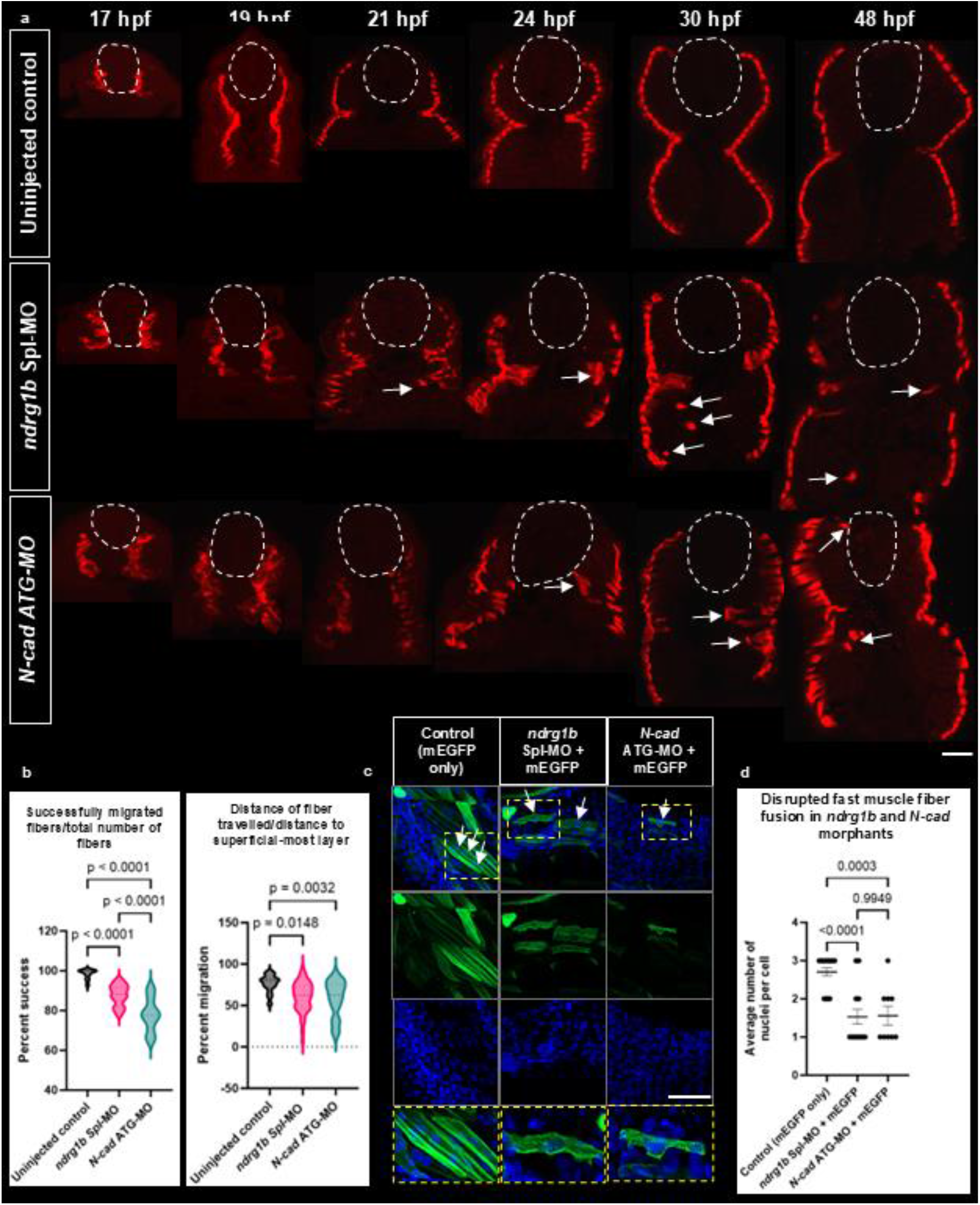
Loss of *ndrg1b* causes defects in slow muscle migration and fast muscle fusion. (a) Slow muscle fibers migrate collectively outward from the notochord and are just deep to the epidermis in control embryos (top). Transverse sections reveal individual fibers that have failed to migrate outward in *ndrg1b* (a, middle) and *N-cad* morphants (a, bottom). Dorsal views of slow muscle fiber migration shown in Supplemental Information (SI8). (b) *ndrg1b* and *N-cad* morphants show a reduced percentage of successfully migrated fibers (left, partially migrated fibers/total fibers x 100) Success rates: control = 98.28%, *ndrg1b* morphants = 87.67%, *N-cad* morphants = 77.95%. Of the fibers that failed to migrate, their “percent migration” is represented by the distance of the individual fiber/distance to the slow muscle layer (right) Distance percent: control = 76.37%, *ndrg1b* morphants = 60.30%, *N-cad* morphant = 58.2%. (c) After slow muscle migration is complete, secondary myogenesis begins in order to increase muscle size and mass. This is marked by the fusion of fast muscle cells into multinucleated fibers, which is N-cad-dependent. The organization of fibers at 48 hpf is severely disrupted in *ndrg1b* and *N-cad* morphants, marked by shorter, undulated fibers with fewer nuclei (wholemount imaging from laterally-positioned embryos). (d) *ndrg1b* and *N-cad* morphants show a reduction in the average number of nuclei per fiber. Control embryos: average = 2.7 nuclei/fiber, 20 cells counted from 5 embryos; ndrg1b morphants: average = 1.53 nuclei/fiber, 17 cells counted from 6 embryos; N-cad morphants: average = 1.55 nuclei/fiber, 9 cells counted from 5 embryos. Ordinary one way ANOVA used for statistical analyses. Bars represent standard error of the mean (SEM). Scale: white = 20 µm.

### Ndrg1b regulates fast muscle fiber fusion

Another aspect of muscle development that is dependent on N-cad function is fast muscle fiber fusion (Hromowyk et al., 2020). After slow muscle migration is complete, the next phase of myogenesis begins, during which fast muscle cells fuse into multinucleated fibers to maximize protein synthesis and promote muscle growth. The fusion of muscle fibers also requires the tight regulation of cell adhesion as cells must properly recognize, attach, and merge membranes with a neighboring cell (Hromowyk et al., 2020). This fusion is manifested by the increased number of nuclei within an individual fiber. Given that slow muscle migration and cell adhesion are disrupted in *ndrg1b* and *N-cad* morphants, we speculated that there would be disruptions in fast muscle fusion as well.

To test this, embryos were injected with mEGFP (as described above), stained with DAPI, and imaged laterally to visualize whole, individual muscle fibers. In uninjected controls, fast muscles successfully fuse by 48 hpf and contain, on average, 2.7 nuclei per fiber (Fig. 5b,c). In *ndrg1b* and *N-cad* morphants, there are clear defects in muscle fiber fusion, which is noted by the disorganization of fast muscle cells (shorter, undulated fibers) and a significant decrease in the number of nuclei per fiber (1.5 nuclei/fiber, on average, for both; Fig. 5b,c).

Interestingly, morphological defects in muscle fibers appear more severe in *N-cad* morphants with shorter and thinner fibers, though this may be attributed to a complete loss of N-cad protein compared to only a partial loss observed in *ndrg1b* morphants. This disruption to fast muscle fiber fusion further implicates Ndrg1b in muscle development, likely through its regulation of N-cad subcellular localization.

Overall, we observe that *ndrg1b* morphants display defects in slow muscle migration and fast muscle fiber fusion, consistent with those reported with the loss of N-cad and confirmed in *N-cad* morphants. These results carry significant implications for understanding the pathology of Charcot-Marie-Tooth Disease Type 4D, a sensorimotor neuropathy which has been linked to mutations in NDRG1 and is characterized by peripheral nerve demyelination and muscle atrophy (Echaniz-Laguna et al., 2007). While NDRG1 has been studied for its role in myelin production (Pietiäinen et al., 2013), this study may be the first to link the etiology of the disease directly to muscle defects through NDRG dysfunction as well.

### Ndrg1b promotes cell adhesion *in vitro*

The work performed above supports a role for Ndrg1b in regulating the intracellular trafficking of N-cad and suggests that the proper subcellular localization of N-cad is required for muscle cell development in an Ndrg1b-dependent manner. While these studies demonstrate a requirement for Ndrg1b in regulating N-cad in muscle cells, they do not quantitatively assay for a direct loss in cell adhesion. To do so, we used a cell dissociation and aggregation assay in which blastula-stage embryos are manually dissociated, plated onto fibronectin-coated dishes, and imaged over several timepoints as cells reaggregate into clusters (Fig. 6a, modified from Song et al., 2013 and Debruyne et al., 2014). Cell aggregation assays have typically been used as a direct test for the functional integrity of E-cadherin-catenin complexes and therefore as a tool to study invasive and non-invasive cell types in cancer (Debruyne et al., 2014). Notably, uninjected control cells aggregate into significantly larger clusters compared to their morphant counterparts (Fig. 6b-d; Note that, as cells are actively dividing, the absolute size of the cell cluster does not increase proportionally with elapsed time and therefore is only compared between groups at individual timepoints). These results strongly support our claim that Ndrg1b is a regulator of cell adhesion through its control of N-cad subcellular localization. Moreover, these findings point to a role for Ndrg1b in very early stages of development, which is perhaps not surprising given its ubiquitous expression throughout development (Fig. 1f), but are the first to our knowledge to position NDRGs at the forefront of intracellular trafficking during embryonic development.

**Figure 6.**
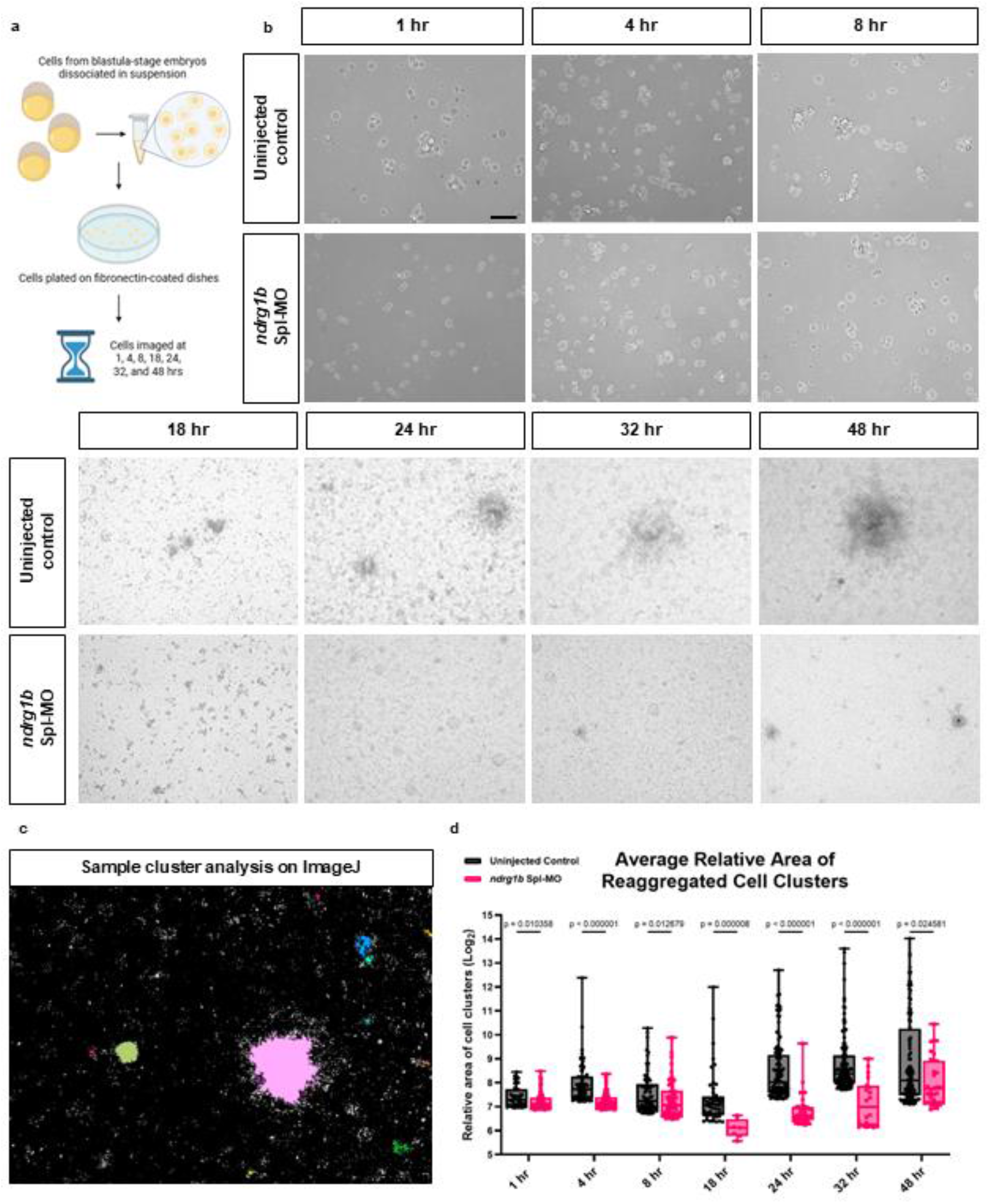
Ndrg1b is required for cell aggregation *in vitro*. (a) Experimental design. Blastula-stage embryos of *ndrg1b* morphants or uninjected controls were deyolked and dissociated at blastula-stage, plated onto fibronectin-coated dishes, and imaged at several timepoints as cells reaggregated. Generated in Biorender. (b) Uninjected control cells re-aggregated in larger clusters over time compared to *ndrg1b* morphant cells. (c,d) ImageJ was used to identify and quantify the relative area of cell clusters. To prevent averaging effects from fewer large clusters and several small clusters, area values obtained from ImageJ were sorted in descending order and the top 5% of values were used in quantifying cluster area. Average relative areas: Controls (in order of timepoints) = 179.97, 338.36, 241.44, 276.68, 620.12, 771.90, 1250.68); Morphants = 157.12, 156.23, 185.44, 66.48, 130.73, 176.77, 344.87).Total number of clusters included in data set for each respective timepoint: AB – 40, 73, 66, 49, 96, 132, 123; 1b MO – 60, 69, 57, 8, 38, 20, 32. Mann-Whitney U test used for statistical analysis. Bars represent standard error of the mean (SEM). Scale: black = 50 µm.

## Discussion

### Summary of findings

In this work we show that *ndrg1b* is ubiquitously expressed throughout early development, consistent with the distribution of *N-cad*, and that the loss-of-function phenotypes in *ndrg1b* and *N-cad* are strikingly similar. We demonstrate that Ndrg1b regulates the trafficking of N-cad, as evidenced by its altered subcellular distribution and accumulation in early and recycling endosomes upon *ndrg1b* loss. These *in vivo* findings are supported by prior studies in prostate cancer cells, where NDRG1 regulates cell-cell adhesion via Rab4-mediated recycling of E-cadherin (Kachhap et al., 2007).

Previous studies in zebrafish have shown that cadherin expression becomes spatially restricted after slow muscle migration, with N-cad highest in superficial slow muscle cells and M-cad more abundant in deeper fast muscle fibers (Cortés et al., 2003; Granato et al., 2003). Our data confirm that N-cad is broadly expressed in slow muscle cells but also reveal previously unreported enrichment of N-cad at junctions between fast muscle fibers, where it appears in cable-like structures along muscle fibers (Fig. 3). While M-cad is known as the dominant cadherin in fast muscle, the enriched N-cad we observe at cell junctions may potentially serve to link muscle fibers together to coordinate muscle contractions. In slow muscle cells, we find N-cad enriched along lateral membranes and at cell-cell junctions, supporting its proposed role in maintaining strong attachments to the overlying epidermis (Kudo et al., 2004; Charvet et al., 2011).

Although NDRGs have previously been implicated in various aspects of protein trafficking, the literature on this topic is primarily focused on conditions of cellular stress including cancer (including but not limited to breast, prostate, esophageal, and pancreatic cancers; Schonkeren et al., 2019; Joshi et al., 2022), hypoxia (Park et al., 2022), and Charcot-Marie-Tooth disease (Echaniz-Laguna et al., 2007; Pietiäinen et al., 2013), and has primarily relied on *in vitro* models that remove critical physiological context from other tissues and organs. Only two studies have previously examined the relationship between NDRG1 and adhesion molecules: one in cultured cancer cells (Kachhap et al., 2007), and one in mice under immersion-induced stress (Miyata et al., 2011). By contrast, our work is among the first to examine the role of NDRGs in protein trafficking during embryonic development *in vivo* in the absence of environmental or metabolic stress.

The novelty of our findings lies in establishing Ndrg1b as a developmental regulator of N-cad trafficking, revealing a role for this protein family beyond stress adaptation. N-cad’s ability to influence both tissue morphogenesis and cancer progression suggests that its regulation must be tightly controlled. We propose that Ndrg1b may act as a molecular adapter that modulates cadherin trafficking under both homeostatic and stress conditions – a hypothesis supported by previous reports that phosphorylated NDRG1 is associated with increased adhesion molecule expression under stress (Miyata et al., 2011) and by unpublished work from our lab. By defining Ndrg1b’s role in an unstressed, developing organism, this study advances our understanding of how adhesion and trafficking are coordinated during morphogenesis, and lays the foundation for future work on how NDRGs bridge developmental and disease processes.

### Physiological relevance

The role of Ndrg1b in regulating N-cad trafficking in muscle suggests it may contribute to processes that are critically dependent on cell adhesion in this tissue. Disruption of adhesion molecules can lead to defective muscle fiber formation, disorganized muscle architecture, and weakened connections between muscle cells and the extracellular matrix. Such impairments are a hallmark of various myopathies, which are often associated with compromised muscle function and increased susceptibility to injury (Tews & Goebel, 1995; Figarella-Branger et al., 2003). These effects are particularly detrimental during periods of growth or regeneration, when muscle tissue must rapidly repair or remodel; under these conditions, defects in adhesion and trafficking may outpace the muscle’s intrinsic regenerative capacity, leading to structural abnormalities, muscle weakness, and eventual atrophy (Demonbreun et al., 2015).

Charcot-Marie-Tooth Disease Type 4D (CMT4D), which has been linked to mutations in NDRG1, is characterized by demyelination of motor and sensory nerves leading to muscle weakness, atrophy, and sensory loss, typically beginning in the distal limbs (Echaniz-Laguna et al., 2007). The prevailing model for NDRG1’s role in CMT4D suggests that defects in Schwann cells (the myelinating cells of the peripheral nervous system) are driven by NDRG1’s impaired ability to traffic the low-density lipoprotein receptor (LDLR), thus resulting in impaired intake of cholesterol which constitutes a significant component of lipid-rich myelin (Pietiäinen et al., 2013). However, our work showing the ubiquitous expression of Ndrg1b and a novel role for Ndrg1b in muscle morphogenesis suggests that muscle atrophy in CMT4D may have multiple cellular origins. These data raise the possibility that muscle atrophy in CMT4D is not solely secondary to peripheral nerve defects but may also stem from intrinsic impairments in muscle cell adhesion and growth (Fig. 5), potentially compounded by altered lipid trafficking (unpublished data; Pietiäinen et al., 2013).

This broader perspective on NDRG1 function is consistent with its paradoxical roles in cancer, where it has been reported to act as both a tumor suppressor and oncogene depending on the cellular context. Our findings add to a growing body of evidence that NDRG proteins are multifunctional regulators whose effects extend beyond individual pathways or cell types. Given the essential role of N-cad turnover in diverse developmental processes – including neural crest and lateral line migration -- the regulation of N-cad trafficking by Ndrg1b may have far-reaching implications across organ systems. The ability to study these interactions in a whole-organism context, as done here, offers new insights into how NDRG dysfunction contributes to disease phenotypes and highlights the need to consider both cell-autonomous and non-cell-autonomous mechanisms in interpreting disease pathology

### Potential cues for upstream activation of NDRGs

NDRGs perform diverse functions that are likely governed by post-translational modifications, subcellular localization, and cellular context. Previous work in our lab has shown that Ndrg1a mediates the endocytosis and degradation of the ATP-demanding NKA pump under hypoxia to conserve energy. While we investigate here the normoxic function of Ndrg1b, preliminary work from our lab suggests that Ndrg1b is upregulated when environmental oxygen levels are low and further increases recycling of N-cad to the cell surface (Brewster lab, unpublished data). These findings support a model in which NDRGs act as tunable environmental sensors that can dial protein trafficking in response to metabolic demands.

Both early development and hypoxia – conditions that regulate *ndrg1b* – are metabolically demanding and characterized by high glycolytic activity. In anoxia, cells rely on glycolysis due to lack of oxygen, while in early embryos, glycolysis supports biosynthetic processes essential for embryonic growth and cell differentiation (Mendelsohn & Gitlin, 2008; Mendelsohn et al., 2008). We hypothesize that lactate, a glycolytic byproduct, serves as a shared upstream signal for Ndrg1b activation. This is supported by studies showing that lactate can bind and stabilize NDRG family proteins (Lee et al., 2015; Park et al., 2015) and that the identified lactate-binding sites in NDRG3 are conserved in Ndrg1a and Ndrg1b (Lee et al., 2015; Park et al., 2022).

Although traditionally viewed as a metabolic waste product, lactate is increasingly recognized as a signaling molecule. It can activate cell surface receptors or directly bind to intracellular targets like NDRGs to influence gene expression and cellular behavior (Lee, 2021; Park et al., 2022). During tissue morphogenesis, high cadherin turnover requires tight regulation of adhesion molecules like N-cad. In contrast, in epithelialized, sessile cells, turnover is reduced. Our findings support a model where lactate-bound Ndrg1b enhances N-cad trafficking during periods of rapid developmental or metabolic change to maintain cell-cell adhesion and homeostasis.

This mechanism is especially relevant to muscle during exercise-induced hypoxia, when ATP consumption outpaces its production. In response, cells shift to glycolysis, leading to rapid lactate accumulation (Nishimura et al., 2010; Sakushima et al., 2020). Hypoxia is a known driver of muscle growth, as seen in strength training, where oxygen depletion and sustained ATP use activate several growth pathways. Stabilization of hypoxia-inducible factor (HIF) under hypoxia promotes angiogenesis to increase oxygen and nutrient delivery and activates satellite cells, muscle stem cells responsible for muscle repair and hypertrophy, causing their differentiation into mature muscle fibers and thus leading to increased muscle mass (Pircher et al., 2021).

Lactate accumulation under hypoxia also triggers an increased secretion of insulin-like growth factor 1 (IGF-1) and growth hormone, further enhancing protein synthesis and regeneration to promote muscle growth (Panzhinskiy et al., 2013; Percher et al., 2021). These combined effects make hypoxia a powerful stimulus for muscle growth and regeneration, and therefore likely depend on enhanced cell adhesion, positioning Ndrg1b as a key regulator in this process. In this model, lactate-stabilized Ndrg1b may promote N-cad localization to the plasma membrane, supporting muscle fiber fusion and coordinated contraction – responses essential for muscle growth and performance. This hypothesis is currently under investigation in our lab.

### Future work

While this study focuses on Ndrg1b’s role in muscle tissue, the broader relevance of these findings lies in the emerging model of NDRGs as key regulators of protein trafficking. Whether Ndrg1b influences N-cad trafficking through direct binding or via intermediary effectors remains an important open question. For example, Kachhap et al. (2007) demonstrated that NDRG1 regulates E-cad recycling through its role as a Rab4 effector, without directly binding the cadherin. This raises the possibility that Ndrg1b may similarly regulate N-cad indirectly through trafficking machinery.

To probe these mechanisms, we are developing new experimental tools. The absence of a reliable antibody against Ndrg1b limits our ability to study its interactions biochemically. However, we have generated GFP-tagged Ndrg1b constructs, which are currently being used to identify binding partners, determine subcellular localization, and perform gain-of-function studies. These tools will also be leveraged to assess whether Ndrg1b is sufficient to promote cell adhesion, and whether its loss-of-function phenotypes can be rescued. Preliminary work using these constructs suggests that the mechanism by which Ndrg1b regulates cell adhesion is through direct interaction with β-catenin, consistent with Co-IP studies performed by Miyata et al., 2011. However, it is also possible that Ndrg1b regulates N-cad through ADP ribosylation factor 6 (Arf6), which is an identified binding partner of NDRGI (Go et al., 2021) and has been shown to regulate the endosomal trafficking of N-cad in migrating neurons of mice (Hara et al., 2016). Given the previously reported literature suggesting NDRGs may alter their function in different cell types, further work is needed to elucidate the mechanism by which Ndrg1b regulates N-cad trafficking in muscle.

Although our current work emphasizes muscle, *ndrg1b* is expressed in multiple tissues, including the hatching gland, kidney, pectoral fin bud, yolk syncytial layer, and central nervous system (Fig. 1). This broad expression suggests a wider functional role. Indeed, over 200 binding partners have been identified for NDRG1 in various tissues (Tu et al., 2007), many of which point to roles in trafficking diverse cellular cargo. Our prior work showed that Ndrg1a regulates the Na+/K+-ATPase in the kidney (Park et al., 2022), supporting the idea that NDRGs modulate the localization of multiple transmembrane proteins (Kachhap et al., 2007; Pietiäinen et al., 2013). Similarly, Ndrg1b may regulate cargos beyond N-cad, a possibility supported by unpublished data from our lab implicating Ndrg1b in the yolk syncytial layer and CNS.

The evolutionary conservation of NDRGs across vertebrates places our findings in a broader phylogenetic context, but also raises new questions. Zebrafish possess two NDRG1 paralogs – *ndrg1a* and *ndrg1b* – arising from the teleost-specific genome duplication. Among these, *ndrg1a* shares greater sequence homology with human NDRG1, including a stretch of three tandem 10-amino-acid repeats in the C-terminus that are missing in *ndrg1b*. The crystal structure of NDRG1 reveals that the C-terminal region may interact with lipids and small metal ions, suggesting it could mediate membrane binding or signaling functions (Mustonen et al., 2020). However, specific domains responsible for protein trafficking remain unidentified.

Ongoing work in our lab involves generating domain-deletion and domain-swap constructs to test whether specific functions – particularly those related to protein trafficking and membrane association – can be mapped to distinct structural regions, including the poorly understood intrinsically disordered C-terminus.

## Conclusions

Over the past few decades, the cellular trafficking pathways of adhesion molecules like N-cad have been extensively studied (Kowalczyk & Nanes, 2012). In this pathway and in many other aspects of vesicle transport, new adapter molecules are frequently being discovered that can help fine tune the complexity of cellular systems. While N-cad’s role is well established, this study identifies a novel mechanism of its regulation through Ndrg1b. Given the vast number of processes in which N-cad is involved – cell specification, migration, adhesion, and many others – it is no surprise that the intricate demands of an organism would call for tight, fine-tuned regulation of this protein. We present Ndrg1b as a novel adapter in the N-cad trafficking pathway that modulates cell adhesion during early development. This work is one of the first to explore Ndrg1b’s role under normoxic conditions in a whole organism. Our findings, supported by unpublished data, suggest that Ndrg1b functions as a bridge in regulating protein trafficking during early development and cellular stress. These insights may reveal broader mechanisms of NDRG function, advancing our understanding of their roles in complex cellular processes. By identifying Ndrg1b as a key regulator of cadherin trafficking, this work not only uncovers a novel mechanism of cell adhesion control, but also expands the paradigm of how pseudoenzymes can orchestrate fundamental morphogenetic processes *in vivo*.

## Materials and Methods

### Zebrafish husbandry and maintenance

Wildtype zebrafish (*Danio rerio)* strains (AB) were reared and utilized using protocols approved by the Institutional Animal Care and Use Committee. Animals were maintained in UV-irradiated, 28.5 °C filtered water and kept in 12:12 light-dark cycles (8am/8pm). Male and female fish were separated by a partition in spawning tanks, with the divider removed at the start of the light cycle to initiate mating. Embryos were collected and staged according to previously described methods (Kimmel et al., 1995). The sex of the embryos used is unknown.

### Morpholino microinjections

Morpholinos used were generated by GeneTools LLC and reconstituted to 1 mM. Embryos were injected in the yolk of 1-4 cell stage embryos with 0.3 pmol of morpholino (1 nL of a 300 uM solution). Glass capillaries used for microinjections were calibrated to 1 nL using the diameter of a droplet suspended in halocarbon oil. Sequence-targeting reagents are listed in supplementary materials.

### CRISPR injections

The online bioinformatics tool CHOPCHOP (https://chopchop.cbu.uib.no/) was used to identify CRISPR target sites in *ndrg1b*. Given the large number of Ndrg paralogs in fish, crispr RNAs (crRNAs) targeting an early and late exon were used in order to “cut out” the gene entirely and prevent functional redundancy and the upregulation of paralogs that may be triggered by nonsense-mediated mRNA decay (Yi et al., 2021). crRNAs (obtained from IDT) targeting exon 2 (5’ – 3’) and exon 14 (5’ – 3’) were first complexed with tracr RNA (IDT, Cat #1072534) to form a 3 µM guide RNA (gRNA) solution, which was then mixed with equal volume 1 mg/mL Cas9 protein (IDT, Cat #1081060) to generate the ribonucleoprotein (RNP) solution. RNP solutions for each guide were combined and injected at 1.5 µM in 3 nL into the single cell of fertilized embryos. F0 crispants following injections were used in some experiments as indicated above, while others were raised to adulthood. Founders from the F0 generation were determined by fin-clipped genomic DNA extraction and amplification of the whole gene (Meeker et al., 2007), with successful knockout resulting in a ∼200 bp product resulting from the deletion of most of the gene. Founder crispants containing a large deletion in the gene product have been outcrossed to WT animals to create an F1 heterozygous population, which has been intercrossed to create an F2 generation which will be screened for homozygous mutants. The generation of a stable mutant line is still underway.

### Riboprobe synthesis

Riboprobes for *in situ* hybridization were prepared using PCR amplification from cDNA synthesis from 6, 12, 24, and 48 hpf embryos (sequences in supplementary materials). RNA extraction was performed with the QuickRNA MicroPrep Kit (Zymo Research, Irvine, CA, USA, Cat #R1051) and cDNA synthesis was carried out using the iScript cDNA Synthesis Kit (Bio-Rad, Hercules, CA, USA, Cat #1708890) according to the manufacturers’ instructions. Amplicon was PCR amplified and RNA synthesized with T7 RNA polymerase (Sigma Aldrich, Cat #10881767001). Riboprobe was labeled with DIG RNA labeling mix (Sigma Aldrich, Cat #11277073910; Lot #57127421). RNA was precipitated with lithium chloride (ThermoFisher, Cat #AM9480; Lot #2644593)

### In situ hybridization

Embryos were fixed in 4% paraformaldehyde solution (PFA) at desired stages and processed for wholemount *in situ* hybridization as previously described (Thisse and Thisse, 2014).

### Hybridization chain reaction (HCR)

Probe sets for HCR were designed by and obtained from Molecular Instruments, along with probe hybridization buffer, fluorescent hairpins, wash buffer, and amplification buffer. Embryos were fixed in 4% paraformaldehyde solution (PFA) at desired stages and processed according to the manufacturer’s protocol (Choi et al., 2016; Choi et al., 2018).

### Transverse sectioning

Transverse sections (50 µm) of fixed and labeled embryos were obtained with the Vibratome 1500. Embryos were positioned in 4% low melt agarose blocks prior to sectioning and mounted on glass slides (Cat #12-550-123) in ProLong antifade mountant (Cat #P36961) under glass cover slips (Cat #50-365-603).

### Immunolabeling

Embryos were fixed in 4% paraformaldehyde solution (PFA) and labeled as previously described (Park et al., 2022). Embryos were stained with Phalloidin-AlexaFluor488 (Thermofisher, Cat #A12379, Lot #s 2409090 and 2615834) and DAPI (SigmaAldrich, Cat #D9542) according to manufacturer’s instructions.

Primary antibodies: anti-N-cadherin (1:100, Abcam, Cat #ab211126); anti-EEA1 (1:100, Santa Cruz Biotechnology, Cat #sc-137130); anti-mGFP/mEGFP (1:100, Abcam, Cat #1218). Secondary antibodies: Goat anti-Rabbit IgG (H+L) Highly Cross-Adsorbed Secondary antibody, Alexa Fluor™ 594 (1:200, ThermoFisher, Cat #A-11037); Goat anti-mouse IgG (H+L) Highly cross-adsorbed secondary antibody, Alexa Fluor™ Plus 488 (1:200, ThermoFisher, Cat #A32723), Goat anti-Mouse IgG (H+L) Highly Cross-Adsorbed Secondary Antibody, Alexa Fluor™ 594 (1:200, Cat #A-11032).

### Compound microscopy

Live, dechorionated embryos were immobilized using the sedative tricaine methanesulfonate at 0.001% w/v in embryo medium E3 (5 mM NaCl, 0.33 mM CaCl2, 0.17 mM KCl, 0.33 mM MgSO4). Sedated or fixed embryos were mounted on depression slides in E3 medium. Lateral views of 24 or 48 hpf hpf embryos were imaged at 5X magnification using an Axioskop II compound microscope (Carl Zeiss, Axioscope II).

### Confocal imaging

Transverse sections mounted on slides were imaged using a ZEISS LSM 900. Magnification is specified in respective images. In comparing expression levels between embryos in the same experimental repeat, all specimens were processed using identical settings. 3D reconstructions were generated using Imaris software. CZI images were imported into the arena, a region of interest was selected, and the ROI was rotated along the X-axis for 200 frames.

### Real-time polymerase chain reaction

Embryos were dechorionated and flash frozen in groups of 3 in crushed up dry ice. RNA extraction was performed with the QuickRNA MicroPrep Kit (Zymo Research, Irvine, CA, USA, Cat #R1051) and cDNA synthesis was carried out using the iScript cDNA Synthesis Kit (Bio-Rad, Hercules, CA, USA, Cat #1708890) according to the manufacturer’s instructions. 12.5 ng cDNA was used in a 20 µl qPCR reaction carried out with a CFX96 Touch Real-time qPCR Detection System (Bio-Rad, Hercules, CA, USA) using the SsoAdvanced Universal SYBR Green Supermix (Bio-Rad, Hercules, CA, USA, Cat #172-5271). Biological triplicates were used for each sample along with technical triplicate. *ef1a* was used as a reference gene as previously validated (Le et al., 2021). Primer sequences are listed in supplemental information. GraphPad Prism 9 software (Prism, San Diego, CA, USA) was used for analysis.

### Western blotting

Embryos were dechorionated and deyolked using Ringer’s solution (116 mM NaCl, 2.9 mM KCl, 5.0 mM HEPES, set pH at 7.2) supplemented with 100 mM PMSF and 10 mM EDTA. Deyolked embryos were flash frozen in groups of 15. Embryos were lysed for 1 hour at 4 °C in Pierce RIPA buffer (Thermo Scientific, #89900) with protease inhibitor (Roche, 05892970001). Boiled lysate was quantified using Pierce BCA assay kit (ThermoFisher, Cat #23227) and 1.25 µg protein loaded per well using Laemmli SDS buffer (Biorad, Cat #1610747). Gel was run with tris/glycine/SDS running buffer (National Diagnostics #EC-870) and transferred with tris glycine transfer buffer (#20960006). Primary antibodies incubated overnight (N-cadherin 1:2500, Abcam, #ab211126; COX-IV loading control 1:2500, Abcam, #ab33985) in TBST. HRP-conjugated secondary antibodies incubated overnight (anti-rabbit 1:5000, Cell Signaling Technology, Cat #7074S, Lot #32; anti-mouse 1:5000, Cell Signaling Technology. Cat #7076P2, Lot #38). Blots were developed with BioRad Clarity Western ECL Substrate (Cat #170-5061) and imaged on an Azure 280 gel doc system.

### mEGFP plasmid synthesis and injections

Membrane EGFP was a gift from Amro Hamdoun (Addgene plasmid # 198056; http://n2t.net/addgene:198056; RRID:Addgene 198056). The plasmid was sent as a bacterial stab and was amplified in LB Agar plates with 100 µg/µl Ampicillin. Single colonies were selected and cultures in LB media with the same antibiotic. The plasmid was purified using the NucleoBond Xtra Midi kit (Cat #740410.50; Lot #2209-009) and 200 pg was injected into 1-4 cell stage embryos.

### Cell dissociation and aggregation assay

Blastula-stage embryos from uninjected controls and *ndrg1b* morphants were harvested in groups of 50 and dissociated by trituration with deyolking buffer (55 mM NaCl, 1.8 mM KCl, 1.25 mM NaHCO_3_) in Hank’s Balanced Salt Solution. Blastoderm cells were harvested by centrifugation for 1 min at 500g. Cells were then resuspended in PBS and spun down in two consecutive washes. The cell pellet was resuspended in 750 µl of L-15 medium supplemented with 0.3 mg/ml L-Glutamine and passed through a 40 µm cell strainer. 500 µl of this cell suspension was plated on 35 mm petri dishes coated with fibronectin at a concentration of 1 µg/cm^2^. Cultures were maintained at 28.5 °C and imaged at 10X magnification using an Axioskop II compound microscope (Carl Zeiss, Axioscope II). Cluster sizes were quantified on ImageJ by adjusting threshold values (between 170-200) to enhance contract and subsequently using the Analyze Particles feature. A minimum of 10 microns was used for 1-8 hours and 15 microns for 18-48 hours. To prevent from averaging effects between a few large clusters and several smaller clusters, the top 5% of identified clusters from ImageJ were compared between uninjected control and *ndrg1b* morphant cells.

