## Supplemental Information for "N-Myc downstream regulated gene 1b is a regulator of cell adhesion during early muscle development"

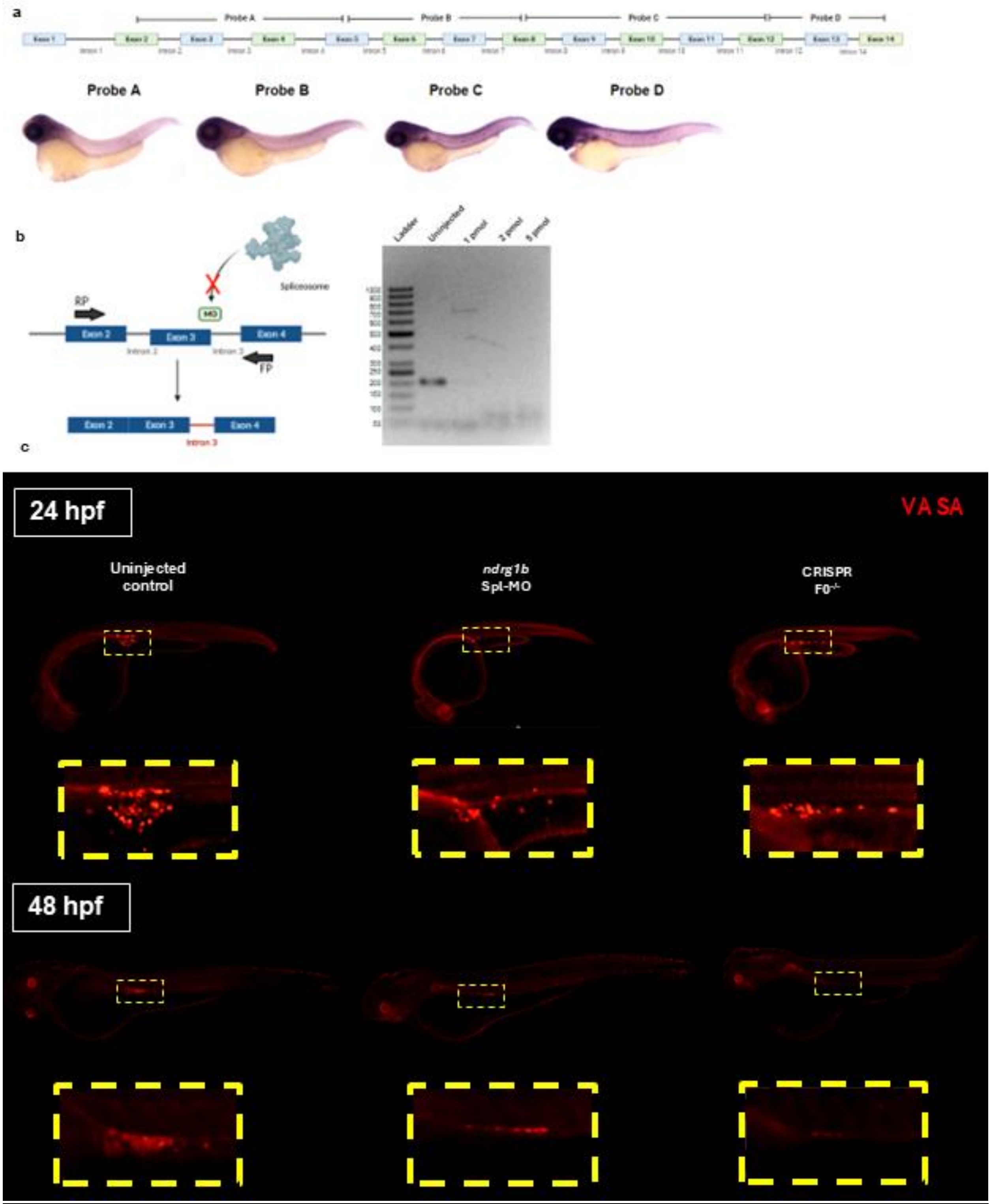


SI1. Tool validation. (a) Comparisons between previously published riboprobe sequences – 932 bp probe from 5' UTR–exon 12 and 458 bp probe from exon 12–3' UTR – have yielded inconsistent results regarding the spatial distribution of *ndrg1b*. To resolve these discrepancies,

we designed and generated four riboprobes of similar lengths (~200-300bp) all within the coding region of *ndrg1b* (supplemental information, probe D used in all subsequent experiments). (b) The splice-blocking morpholino used in experiments was previously used by Li et. al (2016) and was verified using the same methods. The *ndrg1b* splice-blocking morpholino targets a site spanning the exon 3/intron 3 junction and therefore blocks the spliceosome from removing intron 3. This likely results in intron retention and subsequent nonsense-mediated mRNA decay (Li et al., 2016). Efficiency of the MO is tested by primers spanning exon 3 (left, generated in Biorender). In uninjected embryos, mature mRNA, amplification of this region results in a 204 bp product; however no amplicon is detected in splice-blocked embryos. Given that splice-blocking mRNAs do not block maternally inherited mature transcripts, we designed a translation blocking morpholino that binds upstream of the ATG start codon and prevents the binding of the translation initiation complex. However, the lack of commercially available antibodies for Ndrg1b as well as our challenges in generating a custom antibody make this morpholino difficult to validate. The phenocopy between both morpholinos and the ability to validate the splice-blocking morpholino led us to utilize the Spl-MO in our experiments (c) While using CRISPR-knockout embryos would be ideal for the rigor of our work, our lab was unsuccessful in its attempts to generate a stable F2 line of *ndrg1b* crispants. We speculate that this may be due to issues in germline cell proliferation, which have been reported in *ndrg1b* F0 mutants in Medaka (Padilla et al., 2021). Upon investigation, immunostaining against VASA protein (a marker for primordial germ cells), we observe that *ndrg1b* morphants and F0 crispants display reduced number and perhaps defective migration of these cells. These results may underlie our challenges in generating a CRISPR knockout line and further support our use of *ndrg1b* morphants in our subsequent experiments.

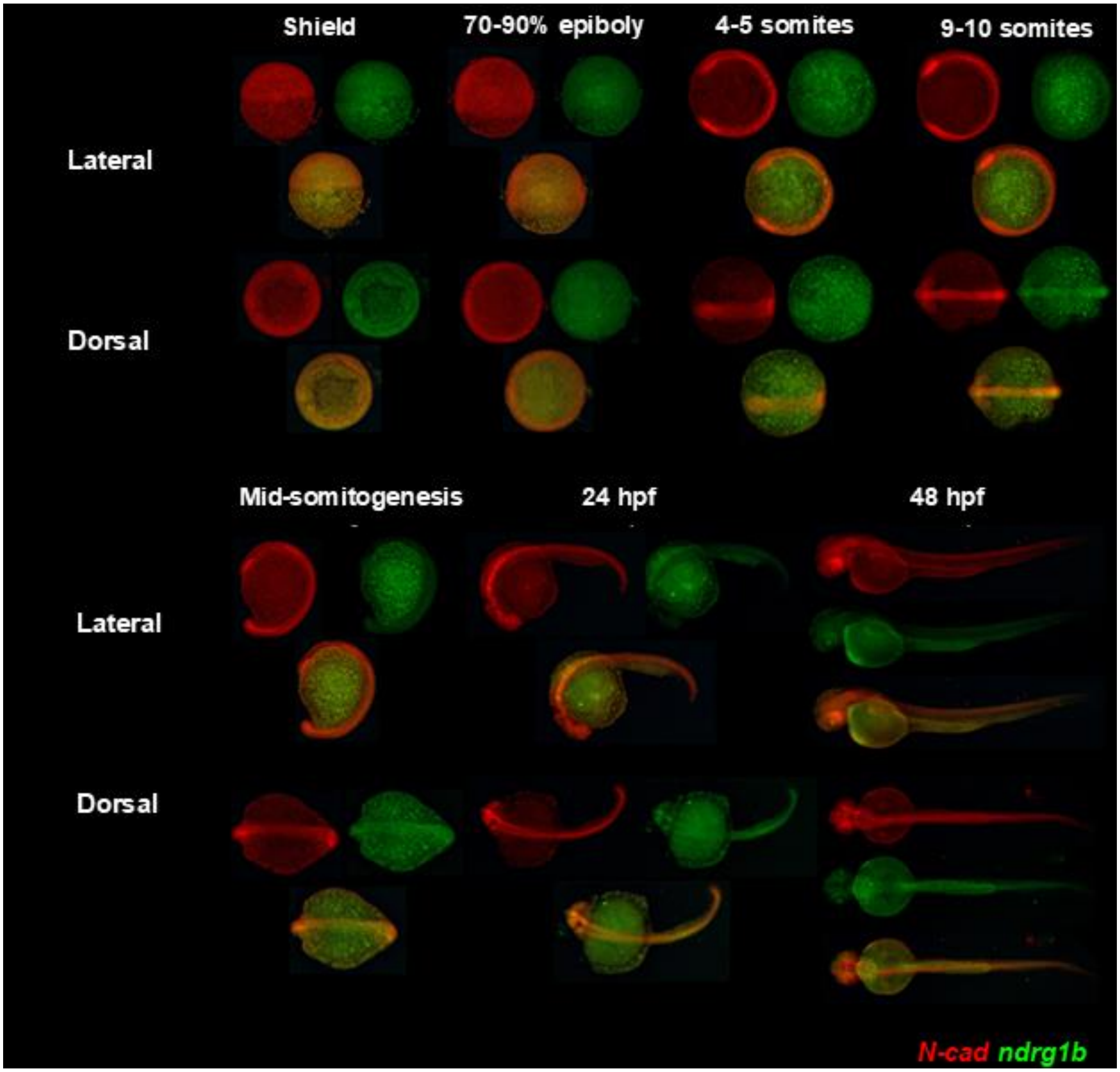

SI2. Lateral and dorsal wholemount views of HCR-FISH double labeling of *ndrg1b* and *N-cad* mRNA. *Ndrg1b* and *N-cad* mRNA colocalize strongly throughout various early stages of development.

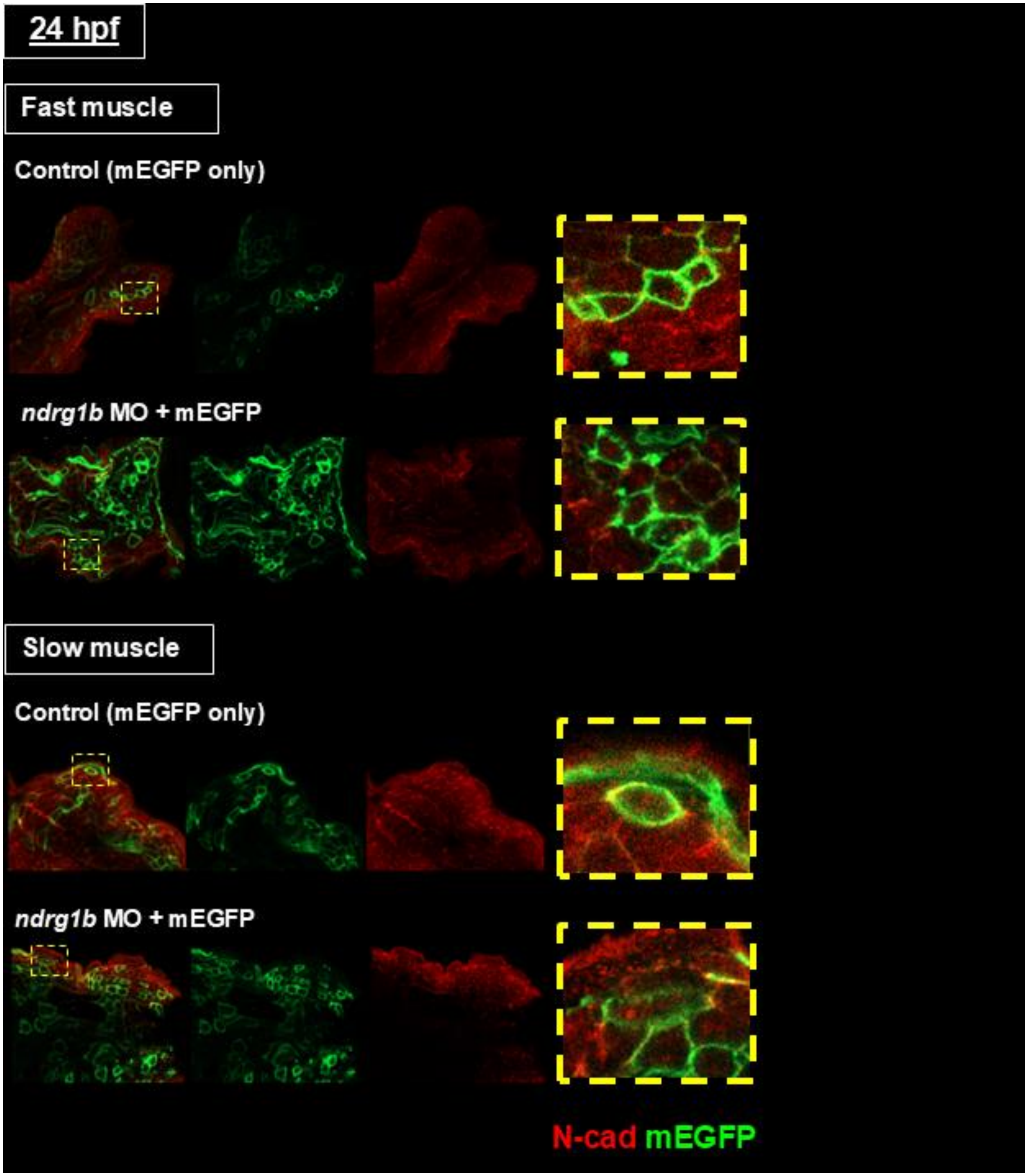


SI3. Individual channels for images shown in Figure 3.

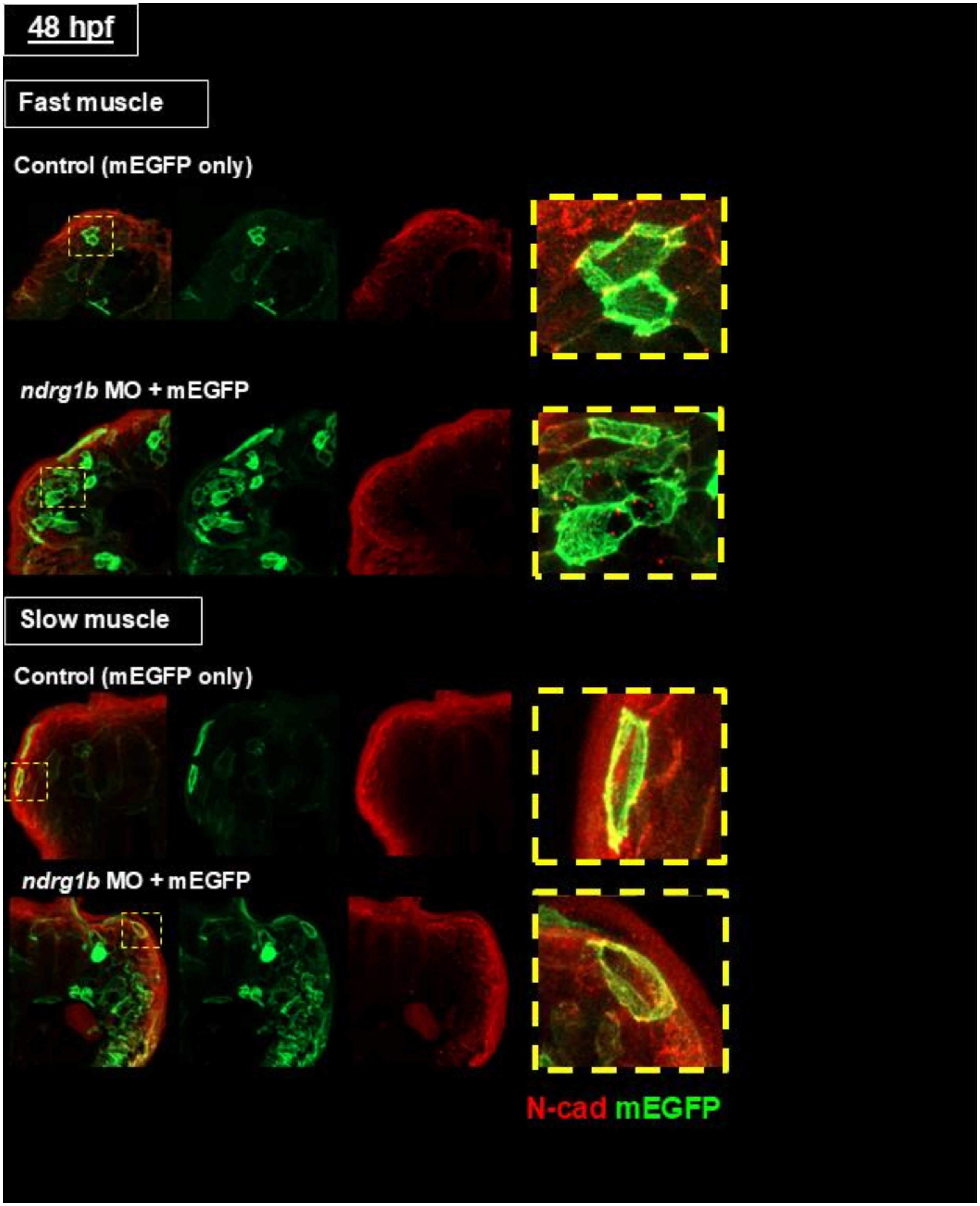


SI4. Individual channels for images shown in Figure 3.

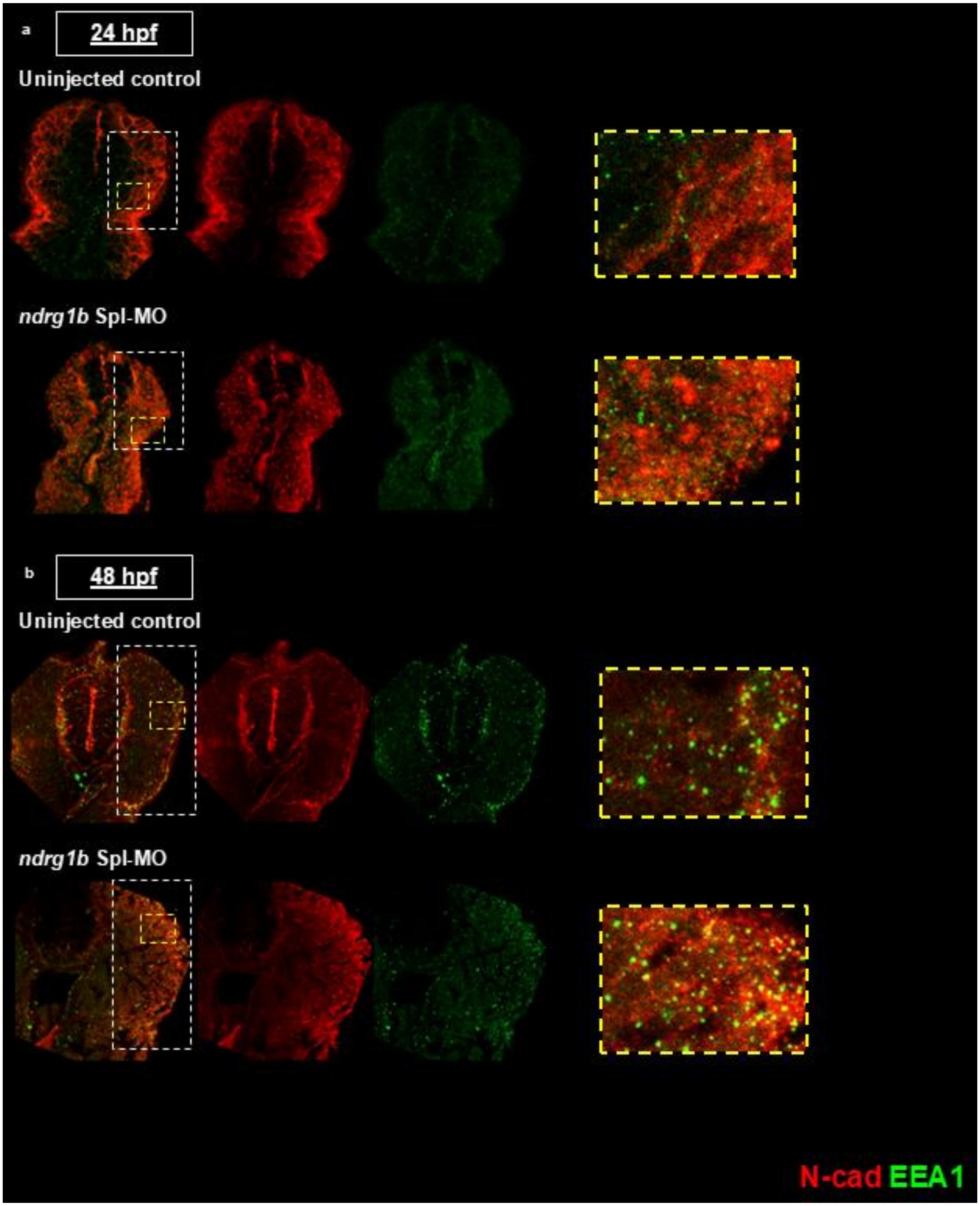

SI5. Individual channels for images shown in Figure 4.

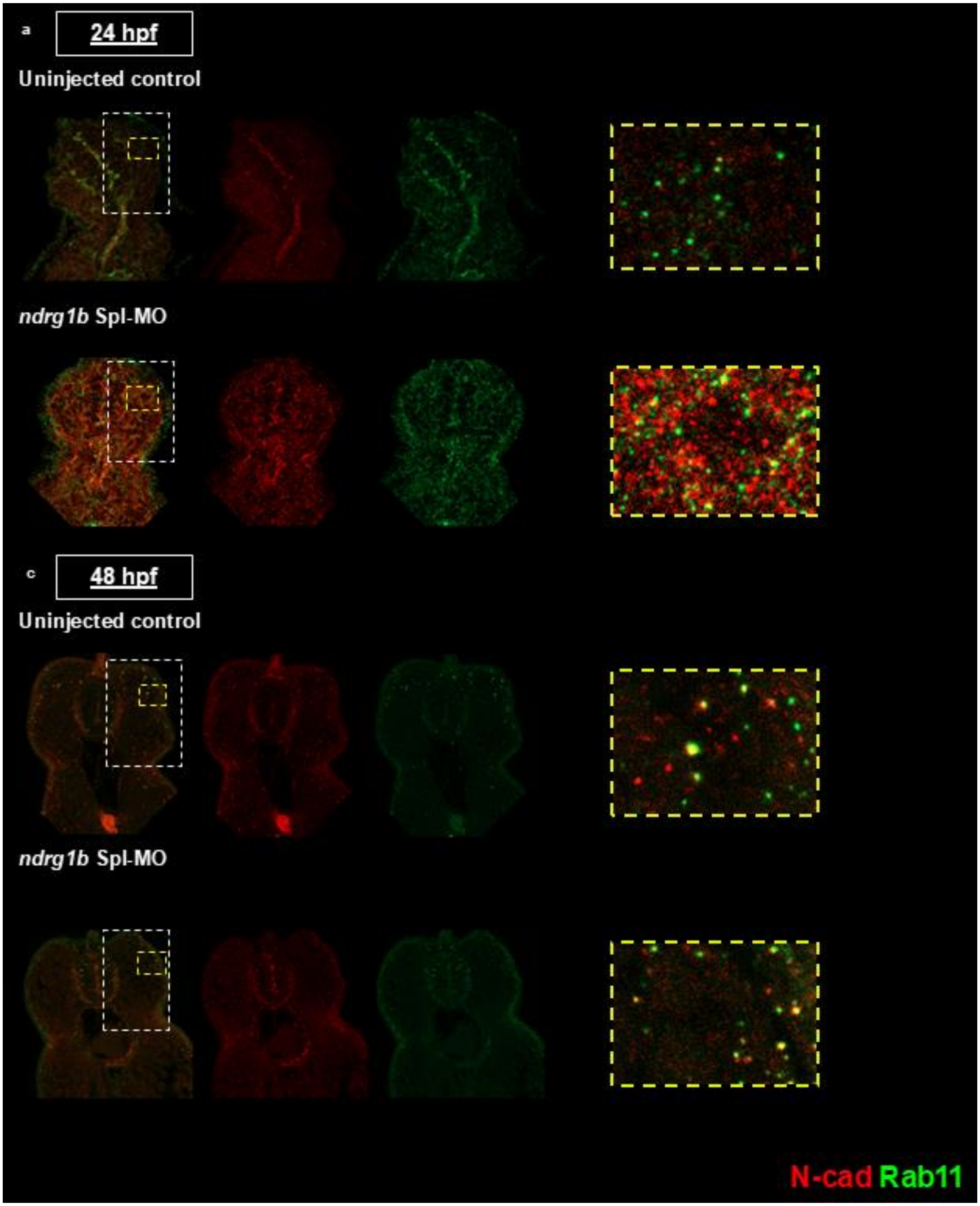

SI6. Individual channels for images shown in Figure 4.

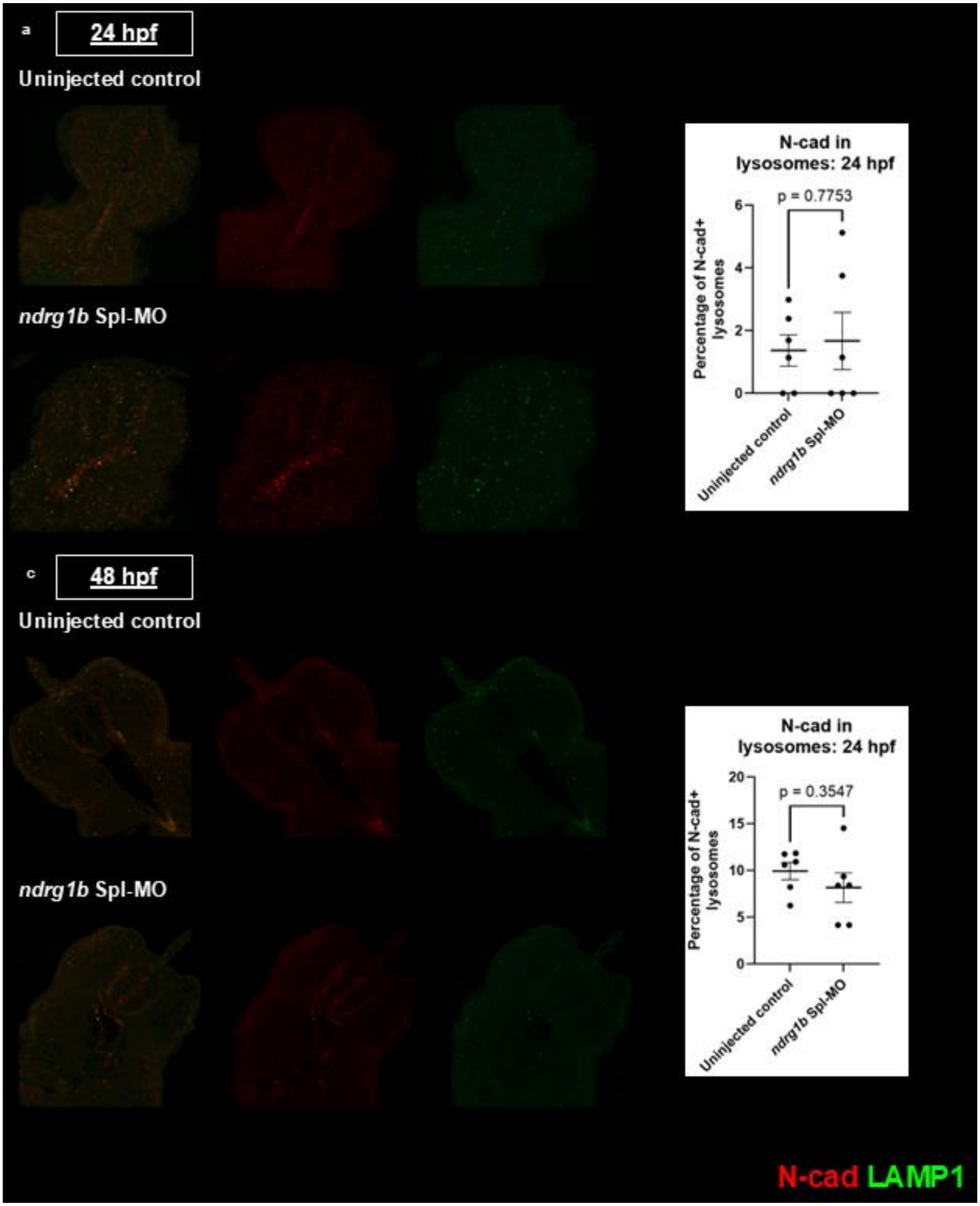


SI7. Double immunostaining for N-cad and LAMP1 (lysosomal associated protein 1). No significant difference was observed in the percentage of N-cad-positive lysosomes between uninjected controls and *ndrg1b* morphants.

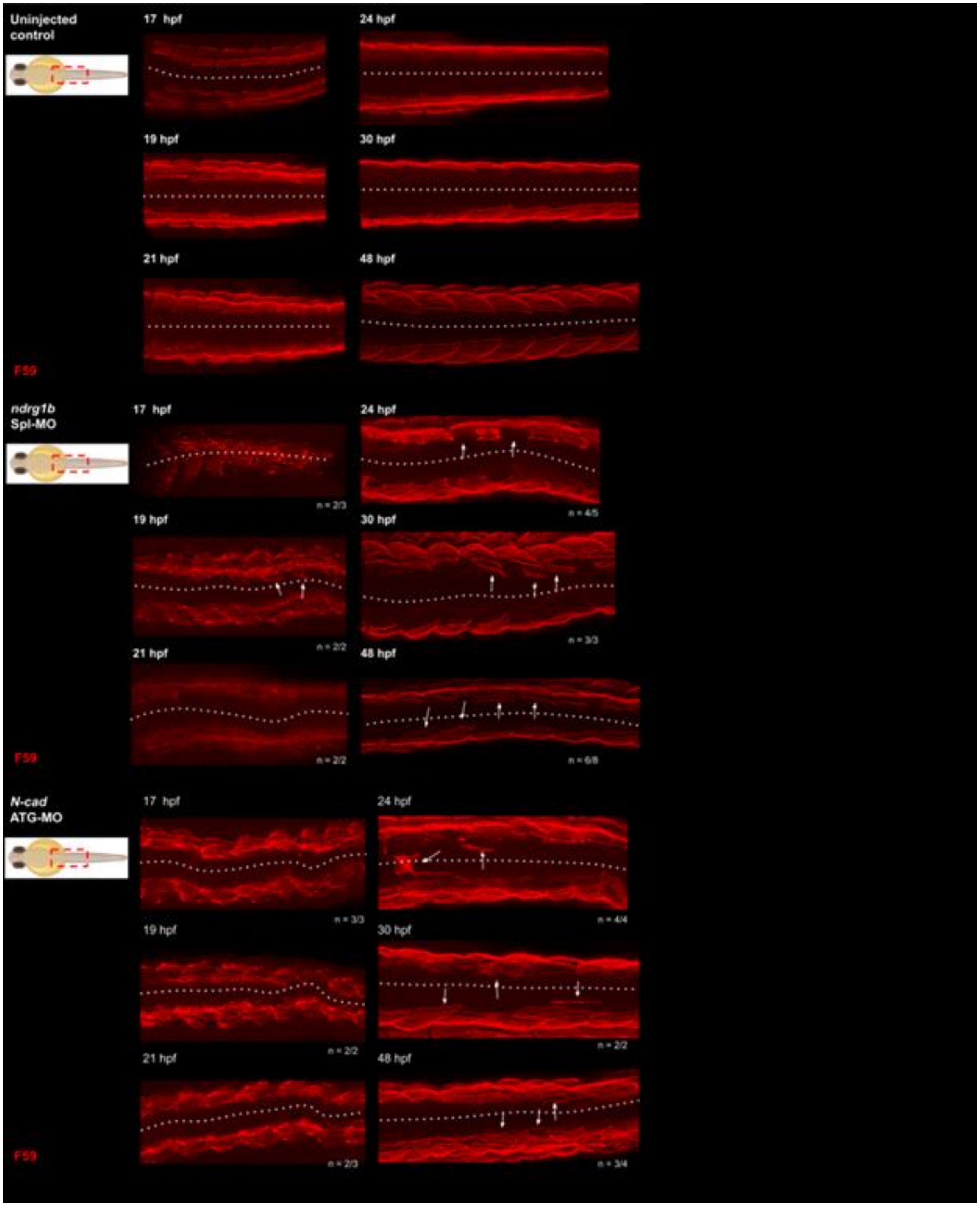


SI8. Dorsal views of slow muscle fiber migration shown in Figure 5.

| Target | Sequence (5' to 3') |
|---|---|
| *ndrg1b* splice-blocking morpholino | AGCATAATGACCCCTGACTCACTCA |
| ndrg1b translation-blocking morpholino | TTGCTGTAAAAGGCTAGTGTGCT |
| CRISPR exon 2 guide sequence | TGTGGAGACCTCCTTTGGAAGGG |
| CRISPR exon 14 guide sequence | TCCTCCAACCGAACCCGCAGCGG |

**Supplemental Table 1. Sequence targeting reagents used in the study.**

| Target | Forward (5' to 3') | Reverse (5' to 3')<br>(T7 promoter underlined) |
|---|---|---|
| *cdh2 (qPCR)* | ACAAGAAGCAGAAGTGTGTGAGC | AGCGTAGGGTCCAGCGTTG |
| *ndrg1b (probe A)* | TTGCACCATGAAGGGTGTCC | TAATACGACTCACTATAGGGCGAGGATATTGGCTCCAG |
| *ndrg1b (probe B)* | CCAATATCCTCGCCCGCTTT | TAATACGACTCACTATAGGACGGAAGGTTGCGATCAGT |
| *ndrg1b (probe C)* | AACTGATCGCAACCTTCCGT | TAATACGACTCACTATACCACCACAGTCAGCCATCTT |
| *ndrg1b (probe D)* | CATGCCCTCTGCCAGTATGA | TAATACGACTCACTATATGCTGAGGCGTGTTTTCAAG |
| Splice-blocking MO validation | CCATCCTGACCTTCCATGAC | GGACCATTGGTAGGCTCTCA |

**Supplemental Table 2. Primer sequences used in the study.**
